# Rearing water microbiomes in white leg shrimp (*Litopenaeus vannamei*) larviculture assemble stochastically and are influenced by the microbiomes of live feed products

**DOI:** 10.1101/2020.08.14.251041

**Authors:** Jasmine Heyse, Ruben Props, Pantipa Kongnuan, Peter De Schryver, Geert Rombaut, Tom Defoirdt, Nico Boon

**Author notes:** Correspondence to: Nico Boon, Ghent University; Faculty of Bioscience Engineering; Centre of Microbial Ecology and Technology (CMET); Coupure Links 653; B-9000 Gent, Belgium; Webpage: www.cmet.ugent.be.

## Abstract

The development of effective management strategies to reduce the occurrence of diseases in aquaculture is hampered by the limited knowledge on the microbial ecology of these systems. In this study, the dynamics and dominant community assembly processes in the rearing water of *Litopenaeus vannamei* larviculture tanks were determined. Additionally, the contribution of peripheral microbiomes, such as those of live and dry feeds, to the rearing water microbiome were quantified. The community assembly in the hatchery rearing water over time was dominated by stochasticity, which explains the observed heterogeneity between replicate cultivations. The community undergoes two shifts that match with the dynamics of the algal abundances in the rearing water. Source tracking analysis revealed that 37% of all bacteria in the hatchery rearing water were either introduced by the live or dry feeds, or during water exchanges. The contribution of the microbiome from the algae was the largest, followed by that of the *Artemia*, the exchange water and the dry feeds. Our findings provide fundamental knowledge on the assembly processes and dynamics of rearing water microbiomes and illustrate the crucial role of these peripheral microbiomes in maintaining health-promoting rearing water microbiomes.

**Originality-Significance Statement:** Most studies on rearing water microbiomes are characterized by sampling resolutions of multiple days and by few replicate cultivations. Through an 18-day sampling campaign in a *Litopenaeus vannamei* hatchery where five replicate cultivations were studied at a sampling resolution of one day, we studied the microbiome dynamics in this system. We show that the community assembly is dominated by stochasticity, which explains the heterogeneity between replicate cultivations. The dynamics of the algal community in the rearing water induced shifts in community composition at two differerent timepoints. Finally, we quantified the contribution of live and dry feed microbiomes to the rearing water community for the first time. We found that the contribution of each source was dependent on its taxonomic composition, the bacterial load caused by the addition of this source and the timing of the introduction. These new insights will aid in the further development of effective microbiome management to reduce the frequency and magnitude of bacterial diseases.

## Introduction

Outbreaks of microbial diseases have posed one of the main impediments to the sustainable growth of the aquaculture industry (Stentiford *et al*., 2017; Shinn *et al*., 2018). Complex changes in the microbial community structure have been hypothesized to be related with disease outbreaks (Xiong, Zhu, and Zhang, 2014; Lemire *et al*., 2015; Dai *et al*., 2020; Huang *et al*., 2020). The aquaculture sector is in need of effective microbial management strategies in order to reduce the occurrence of bacterial diseases. The development and improvement of such strategies is currently hampered by the limited knowledge of the microbial ecology of these systems (De Schryver and Vadstein, 2014; Bentzon-tilia *et al*., 2016).

As compared to terrestrial agriculture, aquatic organisms exist in closer relationship with their surrounding microbiomes (De Schryver and Vadstein, 2014). Numerous molecular studies have found a link between the microbiome of the host and that of the rearing environment (Chen *et al*., 2017; Zheng *et al*., 2017; Sun *et al*., 2019; Angthong *et al*., 2020). The cultivated organisms recrute and enrich specific taxa from their environment (Bakke *et al*., 2015; Yan *et al*., 2016; Li *et al*., 2017; Xiong *et al*., 2019; Zhang *et al*., 2019). For multiple aquatic species, it has been reported that the larvae-associated microbiomes are more similar to the rearing water microbiomes as compared to those in the live or dry feed products (Mcintosh *et al*., 2008; Bakke *et al*., 2013; Giatsis *et al*., 2015). These studies illustrate the importance of the rearing water microbiome in facilitating host-microbiome interactions.

Molecular studies on rearing water microbiomes have shown that the community composition of these systems is dynamic over time (Xiong, Zhu, Wang, *et al*., 2014; Zheng *et al*., 2017; Yan *et al*., 2020), and exhibits large variability across replicate cultivations (Schmidt *et al*., 2017; Chun *et al*., 2018; Li *et al*., 2019; Rita *et al*., 2019; Wiborg *et al*., 2020). The water microbiome experiences frequent disturbances through the addition of live and dry feeds, probiotics and water exchanges, each of which carries with them their own microbiome. It remains largely unknown to what extent the microbial taxa that enter the rearing water can thrive, or even grow (Vadstein *et al*., 2018). The feed products and faecal material produced by the animals cause eutrophication of the water and therefore stimulate bacterial growth (Lucas *et al*., 2010; Chen *et al*., 2017). Zheng et al. (2017) suggested that the shift from one live feed product to another may be responsible for changes observed in the rearing water microbiome, which illustrates the possibility of outgrowth of bacteria from these sources. Live feeds have been associated with potential opportunistic pathogens and antibiotic-resistant bacteria (Mcintosh *et al*., 2008; Hurtado *et al*., 2020; Turgay *et al*., 2020), hence, it is crucial to understand the contribution of these microbes to the rearing water microbiome. Additionally, the magnitude and frequency of disturbances, such as nutrient shocks, have been shown to alter community assembly processes (i.e. stochastic/deterministic balance, Jiang and Patel, 2008; Zhou *et al*., 2014; Santillan *et al*., 2018). Consequently, the regime of these disturbances may be an important driver for the community assembly in the rearing water.

Most studies on the rearing water microbiome are characterized by sampling resolutions of one sample every few days or investigate very few replicate cultivations. Therefore, their resolution does not allow to unveil the complete microbiome dynamics. Additionally, these studies mostly investigate either the bacterial community composition or the bacterial densities, but not both. As has been reported previously, it is crucial to measure both the absolute and relative abundances of microbial taxa to ensure correct interpretation of survey data (Props *et al*., 2017). Also in the context of bacterial disease management in aquaculture, absolute abundances are a crucial factor since virulence of opportunistic pathogens, such as several *Vibrio, Aeromonas* and *Edwardsiella* strains (Henke and Bassler, 2004; Bi *et al*., 2007; Zhang *et al*., 2008), can be density dependent through the regulation of virulence factors by quorum sensing. Additionally, quantification of absolute abundances is important to understand the potential outgrowth of entering micro-organisms, since invader density is an important predictor for invasion success (Jones *et al*., 2017; Kinnunen *et al*., 2018).

To be able to effectively steer the microbiomes towards health-promoting, productive ecosystems, it is imperative to advance our understanding of the assembly, temporal dynamics and drivers of these microbial communities. We performed a sampling campaign on *Litopenaeus vannamei* larviculture and studied the extent to which bacterial taxa present in the rearing water originate from the live and dry feeds or from the exchange water (i.e. sea water that is used to replace part of the rearing water in order to maintain good water quality). We further identified drivers of the community dynamics and determined community assembly processes using an established ecological framework.

## Results

Five replicate *Litopenaeus vannamei* larviculture tanks and all sources that were expected to contribute to the rearing water microbiome, including dry feeds, *Artemia*, algae and exchange water, were monitored over 18 days (life stages N5 to PL10, Figure 1). The rearing water was sampled at a resolution of 3 hours for flow cytometry and once per day for 16S rRNA gene sequencing. Over the cultivation, tank 1 and 4 reached 100% larval mortality at day 13 and 10, respectively (Supplementary Figure 1). Only data from before the mortality event was included for these tanks.

**Figure 1.**
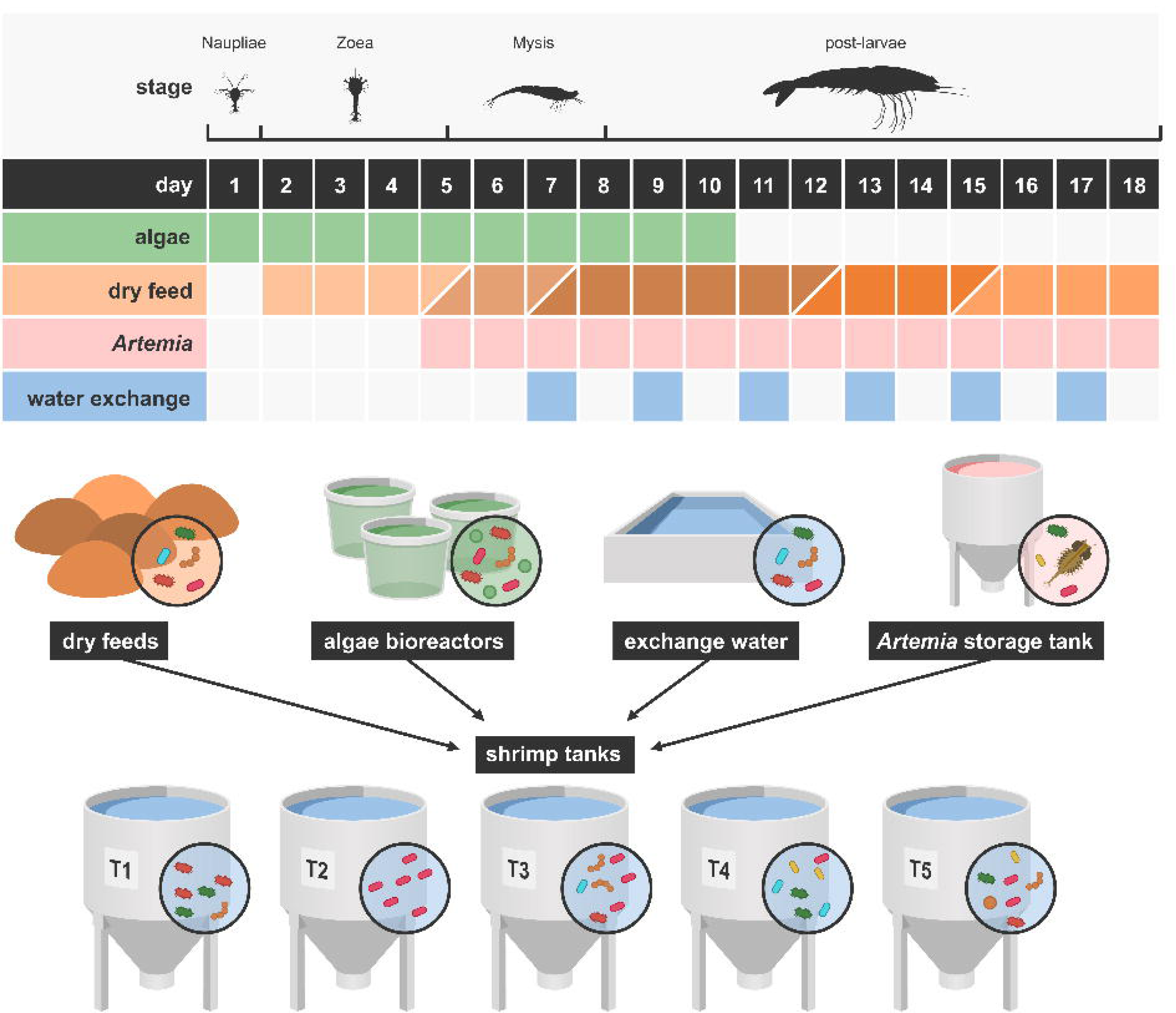
Overview of the experimental setup. Five replicate *Litopenaeus vannamei* larviculture tanks and all sources that were expected to contribute to the rearing water microbiome, including five dry feeds, *Artemia*, algae and exchange water, were monitored over 18 days (life stages N5 to PL10). The upper part of the figure illustrates the timing of addition for each of the external sources to the rearing water. For the dry feeds different shades of brown were used for the different products.

### Bacterial and algal abundances in the rearing water

At the start of the cultivation, after addition of the larvae to the tanks, the bacterial concentration in the water was 4.21 ± 1.44 × 10^5^ cells/mL, and algal cell densities were below the limit of detection (i.e. < 10^3^ cells/mL). Over the following days, a consistent increase in bacterial concentrations was observed for all tanks, reaching an average density of 2.33 ± 0.71 × 10^7^ cells/mL on day 7 (Figure 2A). From day 7 on, the patterns of bacterial cell concentrations started to diverge between the tanks. All tanks experienced a drop in bacterial cell concentrations, followed by a recovery and further increase. The magnitude and timing of this decrease and the ensuing cell growth differed between the tanks and led to differences in bacterial densities of up to 1 log_10_ unit across the tanks.

**Figure 2.**
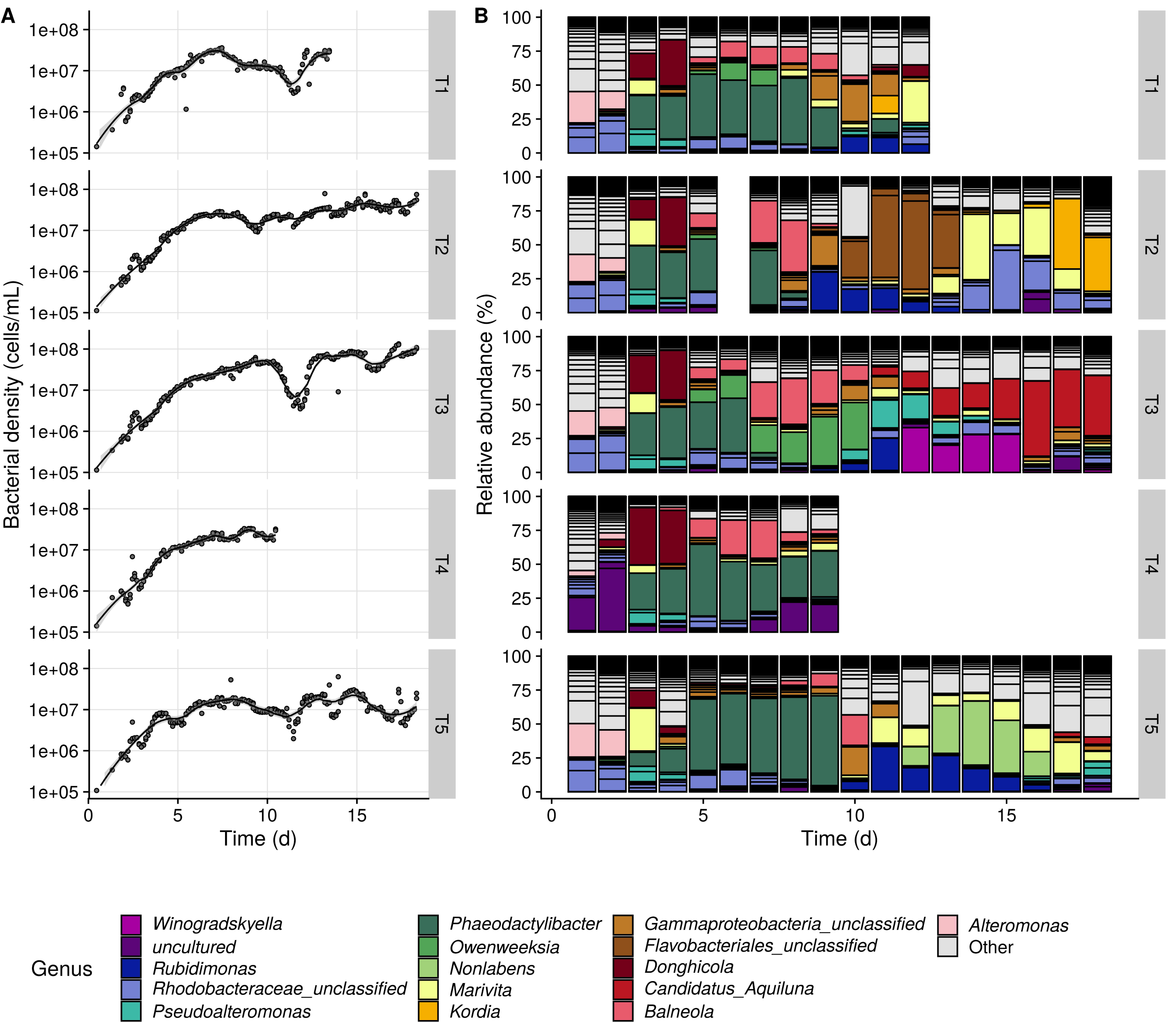
Temporal dynamics of the rearing water bacterial densities (A) and community composition (B) of the five replicate tanks. The OTUs belonging to the 15 most abundant genera are coloured, all other genera were labelled with “Other”. The community composition for tank 2 on day 6 is missing since this sample was not taken.

*Chaetoceros cancitrans* algae were used to feed the larvae over the first 10 days. During this period, algal densities in the rearing water ranged between 1.99 × 10^3^ and 1.03 × 10^5^ cells/mL. After the addition of algae stopped, there was a fast decline in algal densities to below the detection limit on day 12 (Supplementary Figure 2).

### Temporal community dynamics

A PCoA ordination of the Bray-Curtis dissimilarity of the bacterioplankton communities revealed a consistent temporal trend for the replicate tanks (Figure 3 A). A first shift in community composition was observed from day 2 to day 3 (Figure 2 B, Figure 3 A). Beta-diversity partitioning revealed that the observed shift was almost completely (> 99.5 %) attributed to changes in the relative abundances of OTUs that were already present in the system (i.e. ‘abundance variation’), and, thus, not to a large invasion of new taxa (i.e. “turnover”, Supplementary Figure 3). Over this period bacteria were actively growing in the rearing water and bacterial densities increased on average 2.2 fold (Figure 2A). Together, this indicates that the observed community shift was caused by a grow-out that gave rise to an enrichment of specific community members. The OTUs for which there was a sharp increase in relative abundance belonged to the genera *Phaeodactylibacter* (OTU1), *Marivita* (OTU2), *Donghicola* (OTU4) and *Pseudoalteromonas* (OTU13 and OTU27). The initial community composition and dynamics in tank 4 were partly deviating from those of the other tanks (Supplementary Results, Supplementary Figure 4). During the bacterial grow-out from day 2 to day 3, the deviation of tank 4 from the other tanks was reduced, as shown by the lowered average Bray-Curtis dissimilarity between this tank and the other tanks (Figure 3B). The average dissimilarity of tank 5 as compared to the other tanks increased, and this was mainly caused by the more pronounced grow-out of *Marivita* sp. in this tank as compared to the other tanks (Figure 2B, Figure 3B).

**Figure 3.**
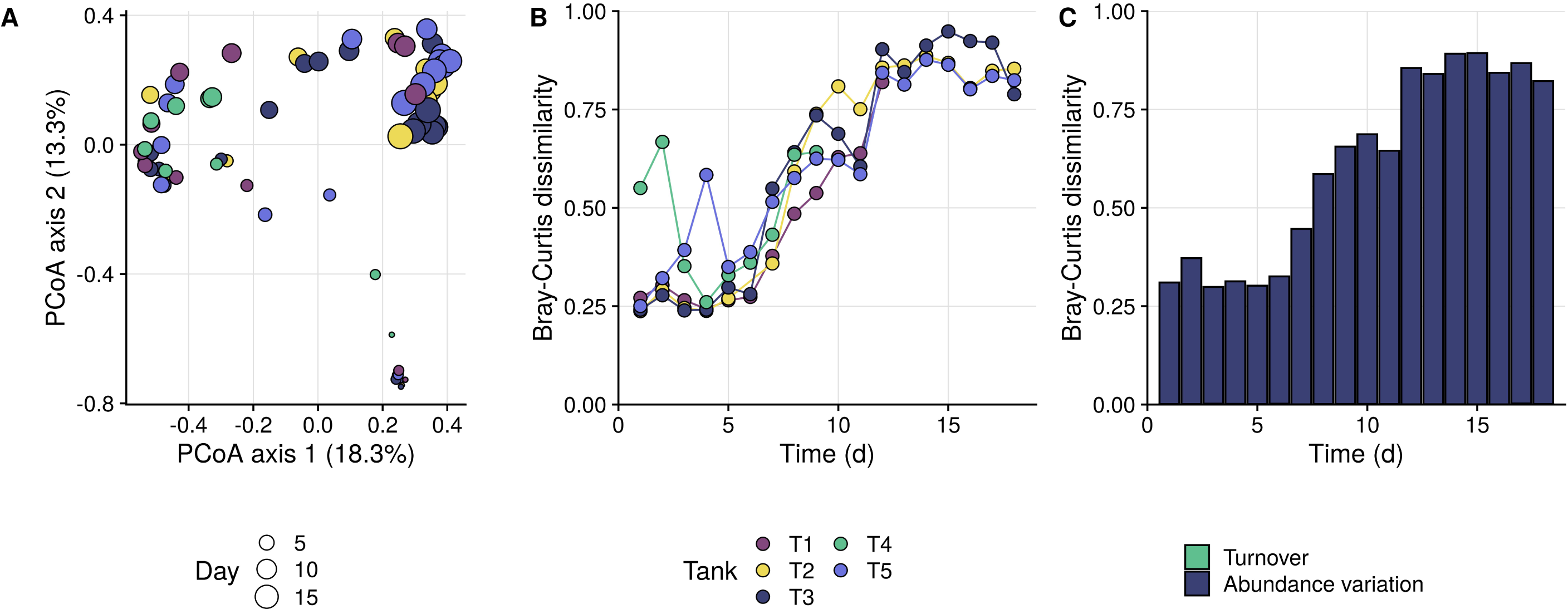
(A) PCoA ordination of the Bray-Curtis dissimilarities of the rearing water microbiomes. Dots are coloured according to the different tanks and the size corresponds to the number of days after the start of the cultivation. (B) Dynamics of the individual Bray-Curtis dissimilarities of each tank compared to the other tanks, per day (e.g. the line of T1 gives the average Bray-Curtis dissimilarities of T1 as compared to the 4 other tanks, per day). (C) Average Bray-Curtis dissimilarities between the 5 tanks, per day, and split according to turnover (i.e. differences in absence/presence of taxa) and abundance variation (i.e. differences in the relative abundances between the tanks). Turnover was responsible for 0.26 % of the dissimilarity on average, and is therefore not visible on the graph.

**Figure 4.**
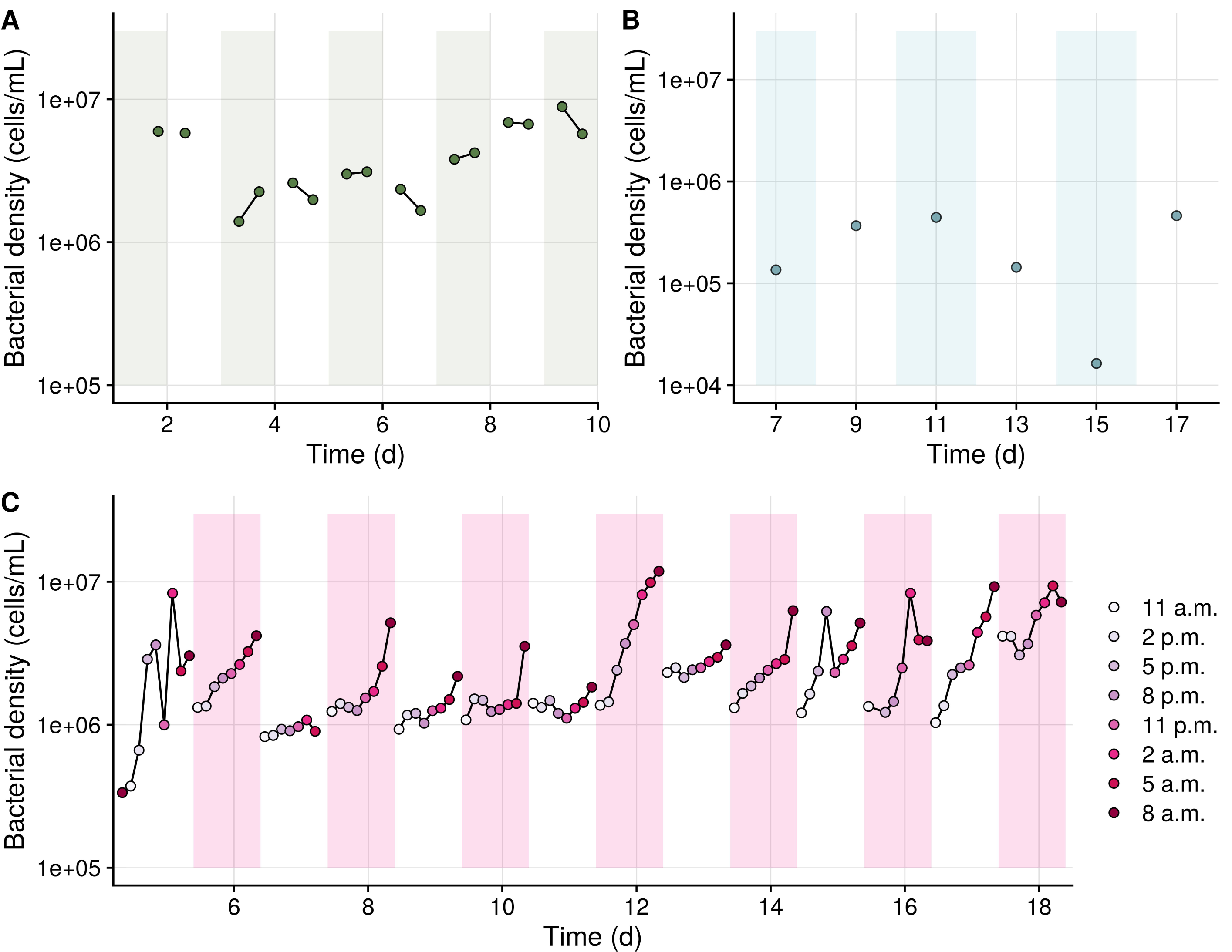
Temporal dynamics of the bacterial abundances in the algal cultures (A), exchange water (B) and *Artemia* storage tanks (C). The different background-colours correspond to the different batches, and the lines connect samples that originated from the same batch. Note that the scale from panel B differs from those of panel A an C.

A second shift established on day 10-11, which corresponded to the final addition of algae as a live feed and the subsequent steep drop in algal abundances in the rearing water on day 11 (Supplementary Figure 2). The absolute abundances of 39 OTUs were significantly (p < 0.001) correlated with the algal densities in the rearing water (Supplementary Table 1). This included some OTUs that had been dominant over the first half of the cultivation, and belonged to the genera *Phaeodactylibacter* (OTU1, r_p_ = 0.55), *Balneola* (OTU5, r_p_ = 0.55), *Owenweeksia* (OTU19, r_p_ = 0.35), unclassified *Saprospiraceae* (OTU30, r_p_ = 0.36) and unclassified *Rhodobacteraceae* (OTU6, r_p_ = 0.43). Along with the steep drop in algal abundance, the relative abundance of these OTUs quickly declined, resulting in the observed community shift. Beta-diversity partitioning confirmed this observation by attributing only a marginal fraction (0.01 - 1.18 %) of the total dissimilarity to turnover and attributing most of it to changes in the relative abundances of resident OTUs (Supplementary Figure 3).

Over time, the dissimilarity between the replicate tanks increased (Figure 3B). Along with the shift in community composition that occurred around day 11, the sharpest increase in the variability between the replicate tanks was observed (i.e. from day 11 to day 12, Figure 3B). Afterwards, the inter-tank variability remained high and the dominant community members differed between the tanks until the end of the cultivation. Beta-diversity partitioning of the dissimilarities between the tanks revealed that the tanks are mainly composed of the same taxa, but they differ from one another due to different relative abundances of these taxa (i.e. 0.26 % turnover on average, Figure 3C).

### Pheripheral microbiomes

Throughout the cultivation, the larvae were fed with different live (algae and *Artemia*) and dry feed products (Figure 1). To be able to evaluate the influence of these peripheral microbiomes on the rearing water microbiome, the cell densities and community composition of these sources were investigated. The microbiomes of the different types of peripheral microbiomes (i.e. algae, *Artemia*, dry feed and exchange water) were different from the rearing water and were significantly different from one another (r^2^ = 0.34, p = 0.001, PERMANOVA; Supplementary Figure 5).

**Figure 5.**
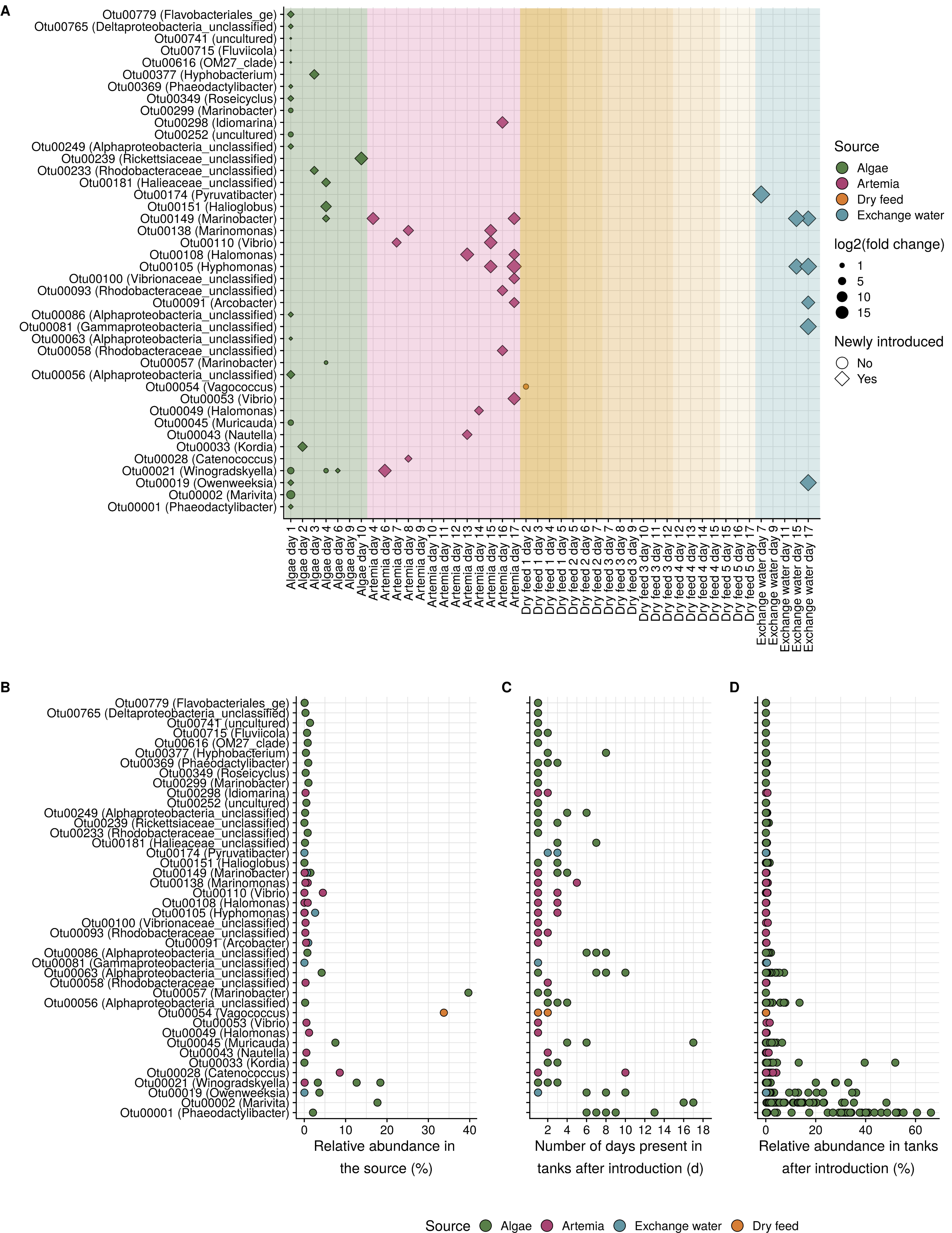
(A) Results of the source tracking analysis based absolute OTU abundances in the rearing water and in the sources. Every dot corresponds to an introduction though one of the pheriperal microbiomes. OTUs that are enriched though the source are indicated with a dot, OTUs that are newly introduced are indicated with a rhombus. The size of the symbol corresponds to the log_2_(fold change) in absolute OTU abundance that was caused on the day before the introduction as compared to the day after the introduction. In case of a newly introduced OTU, the fold change was calculated based on the absolute abundance of the OTU that had been introduced through the source as compared to the day after the introduction. (B) Relative abundance of the introduced OTUs in the source microbiomes from which they originated. An OTU that was introduced multiple times has multiple dots. (C) Number of days the OTU was present in the rearing water microbiomes after introduction. There is a separate dot for each tank, as the residence time of the OTU sometimes differed between the tanks. OTUs that were introduced multiple times have multiple residence times per tank. Dots may overlap. (D) Relative abundance that the OTUs reach in the rearing water after introduction. Every dot corresponds to the relative abundance of this OTU in one tank on one day.

From day 1 (N5) to day 10 (PL2), *Chaetoceros* algae were cultivated in aerated bioreactors and used as live feed. Every day, a single bioreactor was used for two feeding-events. Over the days, bacterial densities in these bioreactors differed up to 1 log_10_ unit (from 1,30 × 10^6^ to 3,28 × 10^7^ cells/mL), and there was a maximum 5 fold-change in bacterial densities between the two feeding-events from the same bioreactor (Figure 4A). The average Bray-Curtis dissimilarity between the communities in the bioreactors was 0.78 ± 0.17, indicating a large batch-to-batch-variability (Supplementary Figure 6). Partitioning of the Bray-Curtis dissimilarity indicated that most variation was explained by differences in relative abundance of the same set of taxa (< 99.5%, Supplementary Table 2). Hence, despite the batch differences, 16 core taxa could be identified (i.e. taxa that were detected in 75% of the samples, Supplementary Table 3).

**Figure 6.**
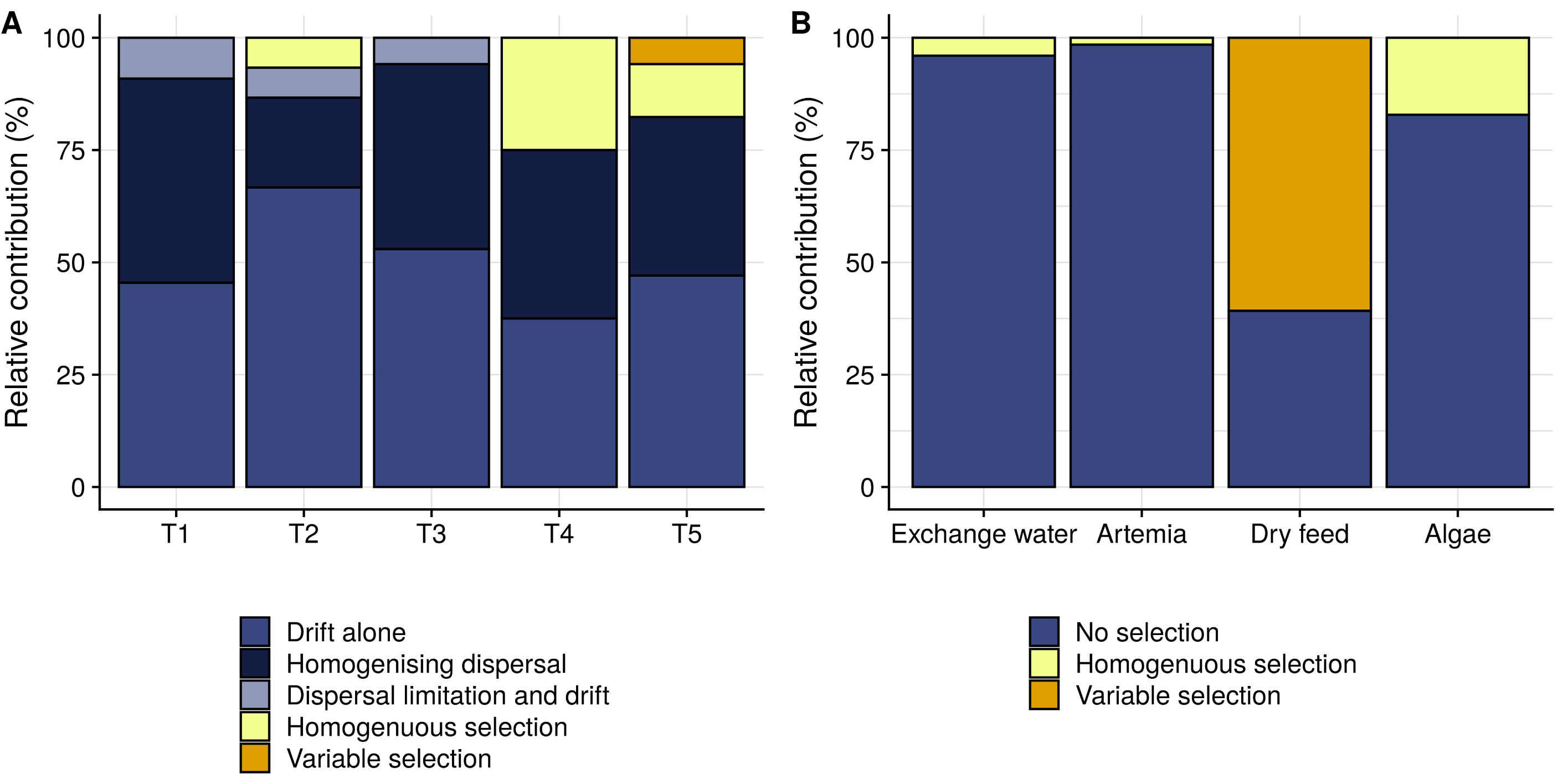
Distribution of the dominant community assembly processes for the individual tanks (A) and from the sources to the rearing water (B), as predicted using the framework of Stegen *et al*. (2013). The blue processes represent the stochastic assembly mechanisms, the yellow processes represent selective mechanisms. Note that both figures have a separate color scale.

From day 5 to day 18 shrimp larvae were fed with *Artemia*, for which a new batch of cysts was hatched every 24 hours and stored at 4°C. The starting bacterial density of the batches ranged from 3.09 × 10^5^ cells/mL to 3.89 × 10^6^ cells/mL (Figure 4C). During the storage, bacterial growth was observed daily, which caused a 1.2 to 10 fold increase in bacterial densities. As was the case for the algae, microbiome composition differed largely between the different batches, as indicated by the average Bray-Curtis dissimilarity of 0.72 ± 0.17, and beta-diversity partitioning revealed that these were mainly related to differences in relative abundances of the taxa that were present (Supplementary Figure 6, Supplementary Table 2). 27 core taxa were identified (Supplementary Table 4).

Five dry feed products were used to feed the larvae throughout the cultivation. Feed product 1 contained a bacterial density of 3.66 × 10^9^ cells/g. For the other dry feeds it was not possible to accurately determine cell densities by flow cytometry (see Supplementary Experimental Procedures). 16S rRNA gene sequencing provided evidence for the presence of a bacterial community also in these feeds, as the detected taxa were distinct from the potential kit contaminants (Supplementary Figure 7). In contrast to the rearing water and other pheripheral microbiomes, the dry feed microbiomes were dominated by Gram-positive taxa (Supplementary Figure 6). There were large differences between the community composition of the different products, which were mainly due to differences in relative abundances of the same set of taxa (Supplementary Table 2).

Water exchanges were performed every other day (30 - 50% of the tank volume) starting at day 7. The cell densities in the exchange water ranged from 1,64 × 10^4^ to 4,62 × 10^5^ cells/mL (Figure 4B). The community composition between the water batches differed with an average Bray-Curtis dissimilarity of 0.83 ± 0.14 (Supplementary Table 2). As for the other sources, these differences were related to differences in the relative abundances of a same set of taxa. Despite the differences between the exchange water on different days, 16 core taxa could be identified (Supplementary Table 5).

### Source tracking

Each source had a different bacterial abundance and was added in a different quantity and with a different frequency. As such, it introduced different microbial loads to the rearing water (Supplementary Figure 8). On average, when algae were added, a bacterial load of 1.22 × 10^4^ cells/mL/d was added to the rearing water. For the *Artemia* this was 1.18 × 10^4^ cells/mL/d, for the dry feed 8.42 × 10^3^ cells/mL/d and for the water exchange 1.45 × 10^2^ cells/mL/d. It should be noted that for the dry feeds this is an underestimation, since we could only determine the bacterial loads accurately for one feed product (see Experimental Procedures).

To investigate to what extent the bacterial taxa detected in the rearing water were associated with the use of the different feed sources, the absolute OTU loads in the rearing water were compared with the OTU load introduced through the feeds and exchange water (see Experimental Procedures for the details on the protocol). This analysis revealed that each of the investigated sources was responsible for the presence of specific taxa in the rearing water. In total, 57 introduction events were detected and the presence of 42 out of the 498 rearing water OTUs could be attributed to the addition of the sources (Figure 5A). Of these OTUs, 26 originated from the algal cultures, 15 from the *Artemia*, 6 from the exchange water and 1 from the dry feed products. Several of the OTUs were associated with multiple sources, either on the same day (e.g. OTU21, *Winogradskyella*) or on different days (e.g. OTU19, *Owenweeksia* sp.).

Remarkably, the introduced OTUs did not belong to the dominant members in the source microbiomes (i.e. only 7 out of 42 OTUs had an abundance > 5% in the source from which they were introduced, Figure 5B). Of the 57 events, 49 were classified as a new introduction, indicating that most of the introduced OTUs were either completely absent in the tank microbiomes, or they had been present before, but were already reduced to below the detection limit.

Several of the introduced OTUs had a long residence time after their introduction, such as OTU2 (*Marivita* sp.) which remained present until the end of the cultivation after its introduction on day 1, whereas other OTUs had residence times of only 1 or 2 days (Figure 5C). The relative abundances of the introduced OTUs in the water could reach up to 60 % (Figure 5D). The initial fold-change that was caused by the introduction of the OTU within 24 hours after its addition was not related to the maximal relative abundance or residence time of this OTU in the rearing water. For example, OTU1 (*Phaeodactylibacter* sp.), which was initially introduced with a relatively small log_2_(fold change) of 1.2, remained present in the rearing water for 6 - 13 days dependent on the tank, and reached relative abundances > 60%. This indicates that even the OTUs that entered the tanks in a relatively low abundance have the potential to grow out to one of the most abundant OTUs in the system. In total, the introduced OTUs represented 37 % of the rearing water community over the entire cultivation.

### Community assembly in the rearing water

The framework developed by Stegen *et al*. (2013) was used to determine dominant community assembly processes in the rearing water over time and to assess the assembly processes that were responsible for the introduction of taxa from the sources. The community assembly in the individual tanks was evaluated through the comparison of the microbiomes on consecutive days (i.e. day *i* to day *i*+1), for each tank separately. For all tanks, the main community drivers were stochastic processes (i.e. 50 % ‘drift acting alone’, 36 % ‘homogenising dispersal’ and 4 % ‘dispersal limitation combined with drift’), and the contribution of selective processes was limited (i.e. 9 % ‘homogeneous selection’ and 1% of ‘variable selection’) (Figure 6A, Supplementary Figure 9).

The assembly process that was responsible for the introduction of taxa from the sources was assessed through comparison of the microbiome in each of the sources to those present in the rearing water one day after the introduction of the source. In this case, only the presence or absence of selective forces was evaluated. In light of the introduction of taxa from the external sources, heterogeneous selection can be interpreted as selection for different properties as compared to those in the source, while homogeneous selection can be interpreted as selection for similar bacteria in the rearing water as compared to those present in the source. The assembly process from the sources differed depending on the source type (Figure 6B). For all sources except for the dry feed, the assembly was dominated by drift (i.e. no selection in favour of bacteria from these sources, nor selection against) with a limited contribution of homogeneous selection (i.e. 4 % for the exchange water, 2 % for the *Artemia* and 17 % for the algae). For the dry feed the assembly was 61 % governed by variable selection.

## Discussion

### Algal populations steer the bacterioplankton dynamics

To obtain microbial control of aquaculture systems it is paramount to understand the drivers of community composition and dynamics (Bentzon-tilia *et al*., 2016). Our data indicated that the communities were continuously changing and that two major shifts occurred (i.e. on day 2 and day 10-11), which were linked to the dynamics of the algal cell densities in the rearing water. The first shift was caused by a rapid grow-out of specific bacterial taxa. There are two possible explanations for this dynamic: an enrichment of bacteria that grow fast on feed and faecal material which accumulates in the tank (De Schryver and Vadstein, 2014; Chen *et al*., 2017) or of bacteria that can grow effectively on algal exudates, given the sharp increase in algal densities (Natrah *et al*., 2014; Mühlenbruch and Grossart, 2018). After this shift the communities were dominated by taxa of which the absolute abundance was significantly correlated with the algal densities (Supplementary Table 1). This supports the hypothesis of outgrowth by algae-associated bacterial taxa. The shift caused an enrichment of the same taxa in all replicate tanks which synchronised the community composition across the tanks (Figure 3B). This can be explained by the community assembly being dominated by homogenous selection and dispersal limitation, two processes that reduce compositional turnover (Supplemantary Figure 9). Algal feeding stopped after ten days which caused a decrease in algal abundance, followed by a decrease in the abundance of algae-associated bacterial taxa. This initiated the second community shift that further increased the divergence in community composition between the replicate tanks.

Phytoplankton is known to steer bacterial communities (Pinhassi *et al*., 2004; Teeling *et al*., 2016; Park *et al*., 2020). Tang et al. (2020) compared cultivations with live and powdered algae and showed that the algae actively modulated the rearing water microbiomes and increased cultivation performance. This principle is widely adopted as the ‘green water technology’, where the presence of algae in rearing water is promoted to improve cultivation performance (Corre et al., 2005; Neori, 2011; Charoonnart and Purton, 2018). The bacterioplankton may be steered by the phytoplankton through different mechanisms. On the one hand, the phytoplankton can compete with bacteria for space and nutrients, and, as such, reduce the outgrowth of bacteria (Mills *et al*., 2008; Fourquez *et al*., 2015). On the other hand, the phytoplankton can interact with and promote specific taxa, as was observed in our study. This interaction can take place through the production of algal exudates that may serve as resources for growth (Smriga *et al*., 2016) or have inhibitory properties (Molina-cárdenas and Sánchez-saavedra, 2017) for specific taxa.

During the cultivation the rearing water is eutrofied which has been shown to increase stochasticity in community assembly and to disturb community stability (Yang *et al*., 2018). Our results indicate that the presence of the phytoplankton community was associated with the presence of specific bacterial taxa and, as such, stabilised the community composition over the replicates. This stabilising effect is in accordance with the study of (Yang *et al*., 2020) which showed that different types of phytoplankton influence the rate at which community change over time, and that this is dependent on the type of algae that were used. However, it should be noted that even though the phytoplankton reduced divergence between the replicate tanks, this did not prevent mass mortality occurring in one of the tanks while algae were abundant in the rearing water (i.e. tank 4). Further research regarding the algae-bacteria interactions in healthy and diseased systems is needed to determine how phytoplankton can be used effectively to steer the rearing water microbiomes towards health-promoting states.

These findings further demonstrate the important role of phytoplankton as a steering factor of aquaculture microbiomes and indicate that management of the stability of the phytoplankton community and its bacterial associates in the rearing water is of great interest during cultivation.

### Rearing water community assembly is dominated by stochasticity

Understanding community assembly processes is imperative to allow the development of effective microbiome management strategies. For example, it can enhance the predictability of factors that determine the establishment of introduced bacteria, such as probiotics (Dini- andreote and Raaijmakers, 2018; Pearson *et al*., 2018), and it may aid in determining optimal process monitoring regimes.

Beta-diversity analysis revealed that the rearing water community composition gradually changes over time (Figure 3A, Supplementary Figure 3), and that through these changes the microbiomes of the replicate tanks increasingly diverged from one another (Figure 3B). The taxa that were present in the replicate tanks were similar, but the relative abundances at which they were present were highly differing (Figure 3C). The community assembly assessment revealed that the temporal dynamics in the rearing water communities are mainly governed by stochastic processes (Figure 6A), which can explain the observed heterogeneity. This stochasticity implies that these larviculture system dynamics are largely unpredictable (Zhou and Ning, 2017) and hence they necessitate continuous monitoring.

Divergence between replicates, such as observed in our study, is a frequently observed phenomenon (Schmidt *et al*., 2017; Chun *et al*., 2018; Li *et al*., 2019; Rita *et al*., 2019; Wiborg *et al*., 2020). Hence, stochastic assembly may be a widespread characteristic aquaculture systems. However, community assembly may depend on the type of rearing system as this determines operational characteristics such as water exchanges/recirculation, feeding frequency, etc. Further research is needed to test the generalisability of our observation.

Not only community composition, but also bacterial densities diverged between the replicate tanks, causing the larvae in replicate tanks to be exposed to different bacterial taxa at different bacterial loads. Given the well-documented link between host and rearing water microbiomes (Zheng *et al*., 2017; Sun *et al*., 2019; Angthong *et al*., 2020), this heterogeneity may have its implications on the reproducibility of cultivation performance. In fact, high mortality and low reproducibility between replicate cultivations are commonly observed in hatcheries (Vestrum *et al*., 2018). However, it should be noted that the cultivated organisms can select for specific taxa from their environment (Yan *et al*., 2016; Li *et al*., 2017; Dai *et al*., 2020), and the exact implications of rearing water community heterogeneity on the reproducibility of cultivation performance remains to be elucidated.

### Peripheral microbiomes are characterised by batch-differences

A hatchery consists of several microbial compartments, including the water column, the larvae, the larval feed, etc. (Goulden *et al*., 2013). Most studies have focussed on the rearing water and host-associated microbiomes, but did not simultaneously investigate the community composition of these peripheral microbiomes. Even though they are recognised as an important factor for biosecurity (Høj *et al*., 2009), limited information is available about batch-variability, bacterial dynamics under storage conditions, etc., in commercial hatcheries.

For each of the peripheral microbiomes a high batch-to-batch variability was observed. These batch-differences were mainly attributed to large differences in the relative abundances of members within a same set of taxa (Supplementary Table 2), and each source was associated with a typical set of “core” taxa (Supplementary Figure 6). Also in terms of bacterial densities, a high variability between and within batches was observed (Figure 4).

The presence of a core set of bacteria in the algal cultures is in accordance with the previously reported specificity of algae-associated microbiomes (Goecke *et al*., 2013; Behringer *et al*., 2018; Fulbright *et al*., 2018; Mönnich *et al*., 2020) (Supplementary Table 3). Many of the taxa that were identified as core members of the algal cultures have been reported to be associated with *Chaetoceros* sp., including *Phaeodactylibacter, Neptuniibacter, Arthrobacter, Marinobacter, Alteromonas, Rhodobacteriaceae, Aestuariibacter, Marinobacter* (Baker *et al*., 2016; Crenn *et al*., 2018; Angthong *et al*., 2020).

For *Artemia*, the taxa that were identified as core members have been previously observed in this system, including *Alteromonas, Vibrio, Nautella, Donghicola, Halomonas* and *Rhodobacteriaceae* (Mcintosh *et al*., 2008; Tkavc *et al*., 2011; Walburn *et al*., 2019; Angthong *et al*., 2020) (Supplementary Table 4). The core microbiome of the *Artemia* storage water harboured several taxa that were classified as *Vibrio* sp., however, due to the limitation of 16S rRNA sequencing we cannot determine wether these bacteria are of concern for the shrimp health in this experiment. This is in accordance with previous studies that reported members of the *Vibrio* genus naturally occur in *Artemia*-associated microbiomes (Lopez-Torres and Lizarraga-Partida, 2001; Thompson *et al*., 2004; Høj *et al*., 2009; Tkavc *et al*., 2011; Interaminense *et al*., 2014). Over the course of 24 hours, bacterial growth which caused up to 1 log_10_-unit differences in bacterial densities was observed. Hence, the storage conditions did not supress bacterial growth. This illustrates that rigorous temperature control is key for bacterial control. The last time point of every batch was sampled for Illumina sequencing, therefore we can make no claims as to which bacterial taxa were growing during the cold storage.

Microbiomes of formulated dry feeds are studied less as compared to those of the live feeds. In our study, the microbiomes of the five dry feed products were dominated by Gram-positive bacteria (**Error! Reference source not found**.), which is in correspondence with a previous report (Giatsis et al., 2015). Some of the families and genera that were detected in the feeds have previously been found in dry feed microbiomes, such as *Bacillaceae* and *Lactobacilli* (Lunestad et al., 2007; Walburn et al., 2019). Feed ingredients, such as processed cell-derived materials, may harbour residual and/or background microbiota and are increasingly used as alternative protein sources for feed production (Cottrell et al., 2020). It is therefore important to note that the bacterial loads measured in this study can comprise viable and/or non-viable cells, and that additional physiological analyses are necessary to verify this. Previous studies have shown these aspects to be affected by environmental parameters as well as on-site handling and usage (Lunestad et al., 2007; O’Keefe and Campabadal, 2015; Walburn et al., 2019). Given the limited research effort towards dry feed microbiomes, further research should elucidate on the functionalities and in situ activity bacteria in dry.

Our data indicated that microbiomes of dry and live feeds as well as exchange water are characterised by large variability, both within and between batches, and hence increased control of their microbiome may contribute to more predictable larviculture production.

### External sources contribute differently to the rearing water microbiome

The rearing water receives frequent bacterial inputs through the addition of live and dry feeds and water exchanges, each of which can contribute to the rearing water microbiome. Several authors have hypothesised that the presence of specific taxa in the rearing water was caused by the addition of these inputs (Zheng *et al*., 2017; Walburn *et al*., 2019; Angthong *et al*., 2020). Nonetheless, the relative importance of each of these inputs for the rearing water community had not yet been quantified (Vadstein *et al*., 2018).

Our source tracking analysis revealed that the microbiomes of all external sources (algae, *Artemia*, dry feeds and exchange water) contributed to the rearing water microbiome, and that ± 10 % of the rearing water OTUs (i.e. 42 out of the 498) were introduced through these sources (Figure 5A). Together, these OTUs were responsible for 37 % of the rearing water community over the entire cultivation. The contribution of the different sources in terms of the number of introduced OTUs, residence time and relative abundances in the rearing water, greatly differed between the sources, with the biggest contribution by the algae, followed by the Artemia, the exchange water and the dry feeds (Figure 5).

The addition of *Artemia* and algae caused similar bacterial loads to the rearing water (Supplementary Figure 8). However, the number of OTUs that were introduced though the algae was higher as compared to those of the *Artemia* (i.e. 26 vs. 15) and the residence times and relative abundances that were obtained by the algae-associated OTUs were higher (Figure 5). This can partly be explained by the fact that the rearing water conditions more frequently favoured the selection of bacteria from the algal cultures as compared to those of the *Artemia* (Figure 6B). Another explanation for the higher contribution of the algae may be that that the algae were added from the start of the cultivation, while the *Artemia* addition only started later (Figure 1). At the start-up, the rearing water was partly disinfected and the bacterial abundance was expected to be below the carrying capacity (De Schryver and Vadstein, 2014). Over the following days feed products and faecal excrements accumulated in the rearing water which caused a gradual eutrophication and bacterial growth (Payne *et al*., 2006), therefore the nutrient availability per cel may have been higher during the introduction of the algae-associated bacteria as compared to those of the *Artemia*. Invasion research has shown that the availability of nutrients is one of the main factors affecting the susceptibility of communities to invasion (Eisenhauer *et al*., 2013; Mallon *et al*., 2015).

Even though cells were detected in one of the dry feeds using flow cytometry, the dry feeds had the lowest overall contribution to the rearing water, which is in accordance with a study of Giatsis *et al*. (2015). This can be explained by a combination of selection against the feed community members (Figure 6B) and the lower bacterial load towards the rearing water as compared to the live feeds. In addition, the dry feed microbiomes may contain a large fraction of non-viable cells, as discussed previously. Despite the large volumetric contribution of the exchange water (i.e. 30 – 50 % of the tank volume), only 6 OTUs were introduced. This can be explained by the low bacterial densities in these exchange waters (2 log_10_-units lower as compared to the live feeds).

Interestingly, the algae were responsible for the introduction of several taxa that correlated with the algal abundances in the rearing water over the first half of the cultivation (i.e. *Phaeodactylibacter* sp., *Marvita* sp. and *Owenweeksia* sp.), and in that way contributed to the observed stabilising effect by the algae-associated bacterial community. On the other hand, the introduced OTUs also included taxonomic groups that are identified as biomarkers for bad shrimp performance (i.e. *Vibrionaceae* sp., originating from the *Artemia*). This illustrates that to maintain health-promoting microbiomes in the rearing water, proper management of the peripheral microbiomes is essential.

Overall, these results illustrate that the peripheral microbiomes have an important contribution to the rearing water microbiome. Given this contribution, careful preparation and storage of these inputs will be paramount to maintain stable, healthy systems. The contribution of each source was dependent on its taxonomic composition, the bacterial load that originated from this source and on the timing of the introduction. Based on this study, the hygiene of the live feed microbiomes should be prioritised as compared to those of dry feeds and exchange water. However, bacterial fluxes can differ dependent on the setup (i.e. cultivation method, disinfection procedures, etc.) (Vadstein *et al*., 2018).

## Conclusions

In this study, we quantified the importance of live and dry feed, and exchange water to the microbial community composition of rearing water in *L*. *vannamei* larviculture. Together these inputs were responsible for 37 % of all bacteria to which the larvae were exposed during the cultivation. The contribution of each source was dependent on its taxonomic composition, the bacterial load caused by the addition of this source and the timing of the introduction. We showed that the temporal community assembly in the rearing water is mainly governed by stochasticity, which corroborates previously documented variable community composition and cultivation performance among replicate cultivations. Additionally, the dynamics of the algal population in the rearing water induced shifts in the bacterial community composition. Our findings provide fundamental knowledge on the sources and assembly processes of the aquaculture microbiome which may aid in the development of effective microbiome management to mitigate bacterial diseases and maintain a health-promoting rearing water environment.

## Experimental procedures

### Rearing of the shrimp larvae

The cultivation of *L*. *vannamei* larvae was performed in five replicate tanks over 18 days (life stages zoea N5 to PL10). The larvae were cultivated indoors, in aerated, 175 L tanks. The five experimental tanks were randomised between other tanks in the facility. Over the course of the cultivation, the larvae were fed every three hours with live and/or dry feed. A range of live and dry feed products were used (Figure 1). From day 5 till day 10, heat-killed (i.e. submerged in boiled water until they are no longer moving) *Artemia* were used as live feed. From day 5 on, live *Artemia* were used. From day 1 to day 10 the larvae were supplemented with *Chaetoceros calcitrans* algae, twice per day. In total, five dry feed products were used, which are labelled as feeds 1 to 5. There was no additional supplementation of antibiotics or commercial probiotics.

Approximately every two days, the physicochemical water quality was assessed tanks (Supplementary Table 6). In order to maintain good water quality, a water exchange between 30 and 50% of the tank volume was performed every other day from day 7 onwards. Larval health was assessed though daily visual inspection (Supplementary Figure 1). Two tanks reached 100% larval mortality during the cultivation (i.e. T4 at day 10 and T1 at day 13).

### Rearing of the live feeds

Algae were cultivated on F2 medium in transparent, aerated bioreactors. Every day a single bioreactor was used to feed the larvae tanks twice (i.e. at 8 a.m. and 5 p.m). Afterwards, the tank was rinsed and cleaned using Sanocare PUR (INVE Aquaculture), according to the manufacturer’s instructions. and refilled to start a new culture.

Every day a new batch of *Artemia* cysts was hatched according to the manufacturer’s instructions. After 20 hours the nauplii were transferred to a refrigerator and stored with aeration at 4°C, to be used as a live feed for the shrimp larvae over the next 24 hours. Every batch was prepared in order to be used from 11 a.m. on the first day until 8 a.m. the following day. Afterwards, the tank was rinsed and cleaned using Sanocare PUR (INVE Aquaculture), according to the manufacturer’s instructions.

### Sampling

Water samples of 1 mL were collected for flow cytometry below the surface of the larviculture tanks before and after every feeding event (i.e. resolution of 3 hours). Since the bacterial load that is added through the feeding is low as compared to the bacterial load that is already present in the tank (i.e. for the algae the daily introduced load is 0.07% of the average bacterial density in the rearing water, and 0.07%, 0.05% and 0.0008% for *Artemia*, dry feed and echange water, respectively), the direct effect of feeding is negligible and the measurement before and after each feeding event can be treated as replicates (i.e. average and median differences in bacterial load before and after feeding are 0.44% and -0.34%, respecively). The water of the *Artemia* storage tanks were sampled at every feeding event, using a sieve to ensure no *Artemia* were included in the sample. The algae bioreactors were sampled, when the algae were added to the cultivation tanks (i.e. twice per day). Samples from the exchange water were taken at every water exchange. All samples for flow cytometry were fixed with 5 µl glutaraldehyde (20%, vol/vol) per mL and stored at 4°C prior to transport. All samples were shipped to Belgium on dry ice for lab analysis.

On day 1 the larvae were added to the water in the evening, and the 1 mL samples for 16S rRNA sequencing that are labelled as “day 1” were taken a few minutes later. For all subsequent days, the sample of the rearing tanks for sequencing was taken every morning, *Artemia* storage and algae bioreactors were sampled for sequencing at the same time. A sample from the exchange water was taken at every water exchange. All dry feeds were samples once. The sequencing samples were stored at -20°C prior to transport. All samples were shipped to Belgium on dry ice for lab analysis.

### Flow cytometry

Prior to FCM analysis, batches of samples were defrosted, acclimated to room temperature and diluted tenfold in sterile, 0.22 µm-filtered Instant Ocean^®^ solution (35 g/L, Aquarium Systems, US). Samples were stained with 1 vol% SYBR^®^ Green I (SG, 100x concentrate in 0.22 µm-filtered DMSO, Invitrogen) and incubated in the dark at 37°C for 20 min. They were analysed immediately after incubation on a BD Accuri C6 Plus cytometer (BD Biosciences, Erembodegem, Belgium) which was equipped with four fluorescence detectors (533/30 nm, 585/40 nm, > 670 nm and 675/25 nm), two scatter detectors and a 20-mW 488-nm laser. Samples were analysed in fixed volume mode (30 µl). The flow cytometer was operated with Milli-Q water (MerckMillipore, Belgium) as sheath fluid and instrument performance was verified daily using CS&T RUO Beads (BD Biosciences, Erembodegem, Belgium). For the feed samples a modified protocol was used (see Supplementary Experimental Procedures).

A subset of 553 samples was measured on-site, immediately after sampling and prior to fixation, on an Accuri C6 (BD Biosciences, Bangkok, Thailand), in order to verify the reliability of the fixation protocol. Pearson correlation between the bacterial and algal densities in samples that were measured fresh (on-site) and fixed (off-site) were 0.9 and 0.9, respectively (Supplementary Figure 11).

### Illumina sequencing

The 1 mL aliquots for DNA extraction were defrosted and pelleted (25 min, 18,200 g, 4 °C). DNA was extracted immediately from the pellet using the ZymoBIOMICS DNA Microprep Kit (Zymo Research, USA). The genomic DNA extract (10 µL) was send out to BaseClear B.V. (Leiden, the Netherlands) for library preparation and sequencing. When algal abundances are high, the 16S sequences originating from chloroplast DNA can make up large portions of the obtained sequences. Because of the presence of the algal populations in part of the samples, PCR clamps were used to block chloroplast amplification and direct the sequencing effort to the bacterial population (Lundberg *et al*., 2013). Amplicon sequencing of the V3–V4 hypervariable region of the 16S rRNA gene was performed on an Illumina MiSeq platform with v3 chemistry, and the primers 341F (5’-CCT ACG GGN GGC WGC AG -3’) and 785Rmod (5’-GAC TAC HVG GGT ATC TAA KCC-3’). Extra samples, including a dilution series of a mock community and a blank, were included as quality controls (see Supplementary Experimental Procedures).

## Data analysis

### Flow cytometry analysis

The flow cytometry data were imported in R (v3.6.3) (R Core Team, 2017) using the flowCore package (v1.52.1) (Hahne *et al*., 2009). The data were transformed using the arcsine hyperbolic function, and the background was removed by manually creating a gate on the primary fluorescent channels. Bacterial and algal populations were separable based on their fluorescence patterns (Supplementary Figure 11). Hamme*s et al* (2008) determined a quantification limit of 100 cells/mL. Since all samples were diluted 10 times, a quantification limit of 10^3^ cells/mL was used.

### 16S rRNA gene amplicon sequencing analysis

Analysis of the amplicon data was performed with MOTHUR (v.1.42.3) (Schloss *et al*., 2009). Contigs were created by merging paired-end reads based on the Phred quality score heuristic and were aligned to the SILVA v123 database. Sequences that did not correspond to the V3–V4 region as well as sequences that contained ambiguous bases or more than 12 homopolymers, were removed. The aligned sequences were filtered and sequencing errors were removed using the pre.cluster command. UCHIME was used to removed chimeras (Edgar *et al*., 2011) and the sequences were clustered in OTUs with 97% similarity with the cluster.split command (average neighbour algorithm). OTUs were subsequently classified using the SILVA v123 database.

The OTU table was further analysed in R (v3.6.3) (R Core Team, 2017). OTU abundances were rescaled by calculating their proportions and multiplying them by the minimum sample size present in the data set. Alpha- and beta-diversities were evaluated using the phyloseq (Mcmurdie and Holmes, 2013) (v1.30.0) and vegan (Oksanen *et al*., 2019) (v2.5-6) packages, respectively. Bray-Curtis dissimilarities were partitioned into turnover and abundance variation, using the betapart package (v1.5.1) (Baselga and Orme, 2012). Differences between groups in the beta diversity analysis were evaluated by means of permutational multivariate ANOVA (PERMANOVA, 999 permutations) of the Bray–Curtis dissimilarity matrix, after confirmation of the homogeneity of the variance in the groups. Core microbiome members were determined using the microbiome package (v1.8.0) (Lahti and Shetty, 2019). The prevalence-threshold that was used for detecting core members was 0.75 (i.e. taxa that were detected in 75% of the samples were considered members of the core). Absolute OTU abundances were calculated based on the bacterial densities as determined though flow cytometry.

### Source tracking

For each day *i*, the absolute abundance of the OTUs in the rearing water of each tank was calculated based on the cell density in the tank on that day, the relative abundance of the OTU in the tank on that day and the tank volume. For each OTU, this absolute abundance is compared to that of the previous day *i*-1. If there is on average an increase from day *i*-1 to day *i* over all tanks, and this increase is observed in at least half of the tanks,this OTU is retained. The criterion is based on minimally half of the tanks in order to allow some biological variability, since microbiomes from replicate aquaculture cultivations are known to be highly variable. For every OTU that was retained, the abundance in the sources that were added to the tank over the course of day *i*-1 to day *i* are verified. If the OTU was present in a specific source, the absolute abundance that was added is calculated based on the cell density in the source that day, the relative abundance of the OTU in the source that day and the added volume of the source If the absolute abundance that was added is greater than the absolute abundance that was already present in the tank on day *i*-1, this source is considered an important contributor of this OTU. Additionally, if the abundance on day *i*-1 was zero, this OTU was tagged as a newly introduced by that source. Since the sampling resolution of the rearing water was 1 sample per day for 16S rRNA gene sequencing, this approach provides a snapshot of the effect of each source within 24 hours after the addition of the source. It should be noted that it is possible that much more OTUs entered the rearing water microbiome through the addition of this source initially, but these already decreased in abundance within 24 hours, and hence, they are not detected. It should be noted that for the *Artemia* cultures the source tracking analysis was performed using the community composition profiles of the *Artemia* storage water and not the *Artemia*-associated microbiome.

### Prediction of community assembly processes

The framework proposed by Stegen *et al*. (2013) was used to determine the dominant community assembly processes in the rearing water. This model relies on phylogenetic and compositional turnover rates to quantitatively estimate influences of drift, selection and dispersal on community assembly, and is explained in detail in the Supplementary Experimental Procedures.

## Data availability

The entire data-analysis pipeline is available as an R Markdown document at https://github.com/jeheyse/SourceTrackingDynamicsShrimp. Raw FCM data and metadata are available on FlowRepository under accession ID FR-FCM-Z2LM (on-site measurements) and ID FR-FCM-Z2LN (off-site measurements). Raw sequence data of the natural and mock communities are available from the NCBI Sequence Read Archive (SRA) under accession ID PRJNA637486.

## Acknowledgements

We would like to thank Phuthongphan Rattayaporn and the ITARC staff for their assistance in the sampling and transport. We would like to thank Funda Torun for feedback on the manuscript and Tim Lacoere for his advice during the DNA extractions and for the design of Figure 1. JH is supported by the Flemish Fund for Scientific research (FWO-Vlaanderen, project 1S80618N). RP is supported by a postdoctoral fellowship of the Flemish Fund for Scientific research (FWO-Vlaanderen). For this project, JH and RP received travel grants from The European Asian aquaculture Technology and innovation Platform (EURASTiP, grant IDs 20180033 and 20180045).

## Contributions

JH, RP, PK, PDS, GR, TD and NB conceived the study. JH, RP and PK performed the sampling campaign. JH and RP performed the lab work. JH analysed the data. JH, RP and NB interpreted the results and wrote the paper. NB supervised the findings of this work. All authors reviewed and approved the manuscript.

