## Supplementary Information for "Rearing water microbiomes in white leg shrimp (*Litopenaeus vannamei*) larviculture assemble stochastically and are influenced by the microbiomes of live feed products"

### for

#### Supplementary Experimental Procedures

##### Flow cytometry protocol for the dry feed products

For the feed samples ±0.1 gram of defrosted feed was added to 1 mL of sterile, 0.22 µm filtered Instant Ocean® solution (35 g/L, Aquarium Systems, US) and prepared using a modified protocol. To release bacterial cells from the dry feed products, these samples were sonicated for 2 min at 10% amplitude using a Q700 sonicator equipped with a cup horn (Qsonica, USA). After sonication the samples were briefly vortexed, and sonicated again for 1 min at 10% amplitude and centrifuged for 3 min at 500 g. The samples were diluted, stained and measured on the cytometer as described above. Finally, the bacterial densities were recalculated per gram of dry feed. Since the sonicated feed products caused a lot of background noise on the flow cytometer, bacterial densities were only quantified for feed product 1, where the population was clearly distinguishable. For the other feed products it was not possible to clearly distinguish between events caused by cells and by background noise (Supplementary Figure 10).

##### Quality controls for Illumina sequencing

The dataset contains samples of a wide range of bacterial densities (i.e. $10^5$ – $10^8$ cells/mL). A dilution series of the ZymoBIOMICS Microbial Community Standard (Zymo Research, USA) was included as a quality control of the DNA-extraction and amplicon sequencing workflow, according to Karstens et al., (2019). The mock consists of 8 bacterial species (*Pseudomonas aeruginosa*, *Escherichia coli*, *Salmonella enterica*, *Lactobacillus fermentum*, *Enterococcus faecalis*, *Staphylococcus aureus*, *Listeria monocytogenes*, and *Bacillus subtilis*) and two fungal species (*Saccharomyces cerevisiae* and *Cryptococcus neoformans*). The mock was serial diluted to bacterial densities of $10^5$, $10^6$, $10^7$ and $10^8$ cells/mL in total volumes of 1mL. The community was diluted in autoclaved PBS that was prepared with microbial DNA-free water (SERVA, SERVA Electrophoresis GmbH, Germany) in an autoclaved and UV-treated Schott bottle. After dilution, the mock communities were pelleted (25 min, 18,200 g, 4 °C), similar to the experimental samples. DNA was extracted from the pellet immediately using the ZymoBIOMICS DNA Microprep Kit (Zymo Research, USA). Based on this control, the expected noise levels in samples with the relevant cell density range is < 5% with a maximal relative abundance of 0.17% and a median relative abundance of 0.005% for the contaminating OTUs. This indicates that contaminating OTUs should have a marginal influence on the results of this sampling campaign. Additionally, an extraction without sample was included in the dataset to assess potential laboratory or reagent contaminants (Supplementary Figure 7). This sample had a DNA concentration below the detection limit and resulted in only 156 reads after processing (as compared to on average DNA concentration of 2.56 ng/μL and 23881 ± 7273 reads for the real samples). Additionally, the microbiome in the blank differed from the microbiomes of all samples included in the dataset, indicating that contamination during extraction and sequencing is negligible.

#### Prediction of community assembly processes

The framework proposed by Stegen et al. (2013) was used to determine the dominant community assembly processes in the rearing water. This model relies on phylogenetic and compositional turnover rates to quantitatively estimate influences of drift, selection and dispersal on community assembly. The framework allows to distinguish two selective processes (i.e. 'homogeneous' and 'heterogeneous selection') and three stochastic processes (i.e. 'drift alone', 'homogenising dispersal' and 'dispersal limitation and drift'). 'Homogeneous selection' is the scenario in which a consistent selective pressure is the primary cause of low compositional turnover. 'Heterogeneous' selection is the scenario in which environmental conditions (biotic or abiotic) change over time or space, which causes a high compositional turnover. Under dominance of stochastic processes, community dispersal will influence the magnitude of compositional turnover. 'Homogenising dispersal' is the scenario under which there is weak selection, but a high dispersal rate (exchange) between a given pair of communities, which homogenises community composition, resulting in low compositional turnover despite the influence of stochasticity. 'Dispersal limitation and drift' is the scenario under which there is weak selection and limited dispersal between communities which can lead to marked ecological drift, and, hence, high compositional turnover.

In a first step, the occurrence of selection is detected based on phylogenetic distances between communities, which are quantified using the $\beta MNTD$ (β-mean nearest taxon distance). $\beta MNTD$ quantifies the phylogenetic distance between each OTU $i$ in one community $k$ and its closest relative $j$ in a second community $m$:

$$\beta MNTD = 0.5 \left[ \sum_{i_k = 1}^{n_k} f_{i_k} \min\left(\Delta_{i_k j_m}\right) + \sum_{i_m = 1}^{n_m} f_{i_m} \min\left(\Delta_{i_m j_k}\right) \right]$$

where $f_{i_k}$ is the relative abundance of OTU $i$ in community $k$, $n_k$is the number of OTUs in community $k$ and $\min\left(\Delta_{i_k j_m}\right)$ is the minimum phylogenetic distance between OTU $i$ in community $k$ and all OTUs $j$ in community $m$. Next, a null distribution of $\beta MNTD$ is calculated under the assumption of absence of selection, by iteratively shuffling species names and abundances across the tips of the phylogeny and recalculating $\beta MNTD$. The observed $\beta MNTD$ values are then normalised based on the average $\beta MNTD_{null}$ and the standard deviation of $\beta MNTD_{null}$:

$$\beta NTI = \frac{\beta MNTD_{obs} - \overline{\beta MNTD_{null}}}{sd(\beta MNTD_{null})}$$

The resulting $\beta NTI$ can be interpreted as the number of standard deviations $\beta MNTD_{obs}$ is diverging from the $\beta MNTD$ that is expected in the absence of drift. Stegen *et al.* (2013) defined $\beta NTI$ values < -2 or > 2 indicative for significantly less than or greater than expected phylogenetic turnover, respectively. $\beta NTI$ > 2 corresponds to communities with highly divergent ecological similarity and is interpreted as an indication of 'variable selection', while $\beta NTI$ < -2 corresponds to communities with high ecological similarity and is interpreted as an indication of ‘homogeneous selection’. A $|\beta NTI| < 2$ indicates that selection is not a major determinant of the community assembly.

In the following step compositional turnover is used to estimate the influence of drift on the community. Hereto the Raup-Crick Bray-Curtis dissimilarity ($RC_{bray}$) is used. The resulting $RC_{bray}$ is a value between 1 and -1, and is a measure that indicates whether two communities share fewer or more taxa than to be expected by random chance. Stegen *et al.* (2013) defined values of $RC_{bray}$ > -0.95 or < 0.95 indicative for significantly less than or greater than expected compositional turnover, respectively. $RC_{bray}$ < -0.95 implies the communities have a higher similarity than expected in case the community assembly is governed by drift. Hence this is indicative for ‘homogenising dispersal’ (i.e. dispersal is high enough to cause low turnover by overruling other assembly processes such as drift). $RC_{bray}$ > 0.95 implies the communities have a lower similarity than expected in case the community assembly is governed by drift alone, and, hence, this is indicative for ‘dispersal limitation’ and the resulting intensified drift is a major determinant for community assembly. A $|RC_{bray}| < 0.95$ indicates that drift alone is the major determinant of the community assembly.

The model was implemented in R (v3.6.3) (R Core Team, 2017) according to https://github.com/stegen/Stegen_etal_ISME_2013, using the ape package (v5.3) (Paradis *et al.*, 2004) to construct a phylogenetic tree and the picante package (v1.8.1) (Kembel *et al.*, 2010) for calculating the mean nearest taxon indices. The number of iterations that was used to determine the null distribution was 999 for $\beta MNTD$ and $RC_{bray}$.

#### Supplementary Results

##### Microbiome characteristics at start-up

The initial community composition and dynamics in tank 4 was deviating from that of the other tanks (Supplementary Figure 4). Before the addition of the larvae to the water, the bacterial density in this tank was slightly higher as compared to that of the other tanks. Approximately 6 hours later, which corresponds to 1 hour after the addition of the larvae, the difference in bacterial abundance became larger and the abundance in tank 4 was almost twice as high of that in the other tanks. By the next day (i.e. day 2), all tanks experienced a 3-fold increase of the microbial densities, except for tank 4, for which a small decrease was observed. In terms of composition, tank 4 contained an abundant OTU that was classified as an uncultured *Alteromonadaceae* (relative abundance of 24.41% and 46.47% on day 1 and 2, respectively). In the other tanks, this OTU was present at a relative abundance < 1% during start-up, and increased to relative abundances of < 5% throughout the cultivation.

#### Supplementary Tables

Supplementary Table 1 – Identity of the OTUs for which there was a significant ($p < 0.001$) Pearson correlation between the absolute OTU and algal abundances. Maximal relative abundance in the rearing water for each OTU is reported as a measure of the relative importance of this OTU in the system. OTUs are sorted according to the maximum relative abundance.

| OTU | Genus | $r_p$ | p-value | Maximal relative abundance (%) |
|---|---|---|---|---|
| OTU1 | *Phaeodactylibacter* | 0,55 | $2,58 \times 10^{-8}$ | 66,17 |
| OTU5 | *Balneola* | 0,55 | $3,87 \times 10^{-8}$ | 38,15 |
| OTU19 | *Owenweeksia* | 0,35 | $9,78 \times 10^{-4}$ | 36,20 |
| OTU30 | Unclassified *Saprospiraceae* | 0,36 | $5,41 \times 10^{-4}$ | 19,62 |
| OTU6 | Unclassified *Rhodobacteraceae* | 0,43 | $3,43 \times 10^{-5}$ | 15,46 |
| OTU111 | *Peredibacter* | 0,41 | $8,63 \times 10^{-5}$ | 8,14 |
| OTU63 | Unclassified *Alphaproteobacteria* | 0,42 | $4,11 \times 10^{-5}$ | 7,34 |
| OTU41 | *Cohaesibacter* | 0,39 | $1,78 \times 10^{-4}$ | 6,98 |
| OTU43 | *Nautella* | 0,43 | $3,03 \times 10^{-5}$ | 6,30 |
| OTU139 | Uncultured *Rubinisphaeraceae* | 0,40 | $1,23 \times 10^{-4}$ | 3,53 |
| OTU98 | *Maricaulis* | 0,37 | $4,81 \times 10^{-4}$ | 3,12 |
| OTU101 | *Candidatus Uhrbacteria* | 0,54 | $8,28 \times 10^{-8}$ | 2,43 |
| OTU81 | Unclassified *Gammaproteobacteria* | 0,39 | $1,72 \times 10^{-4}$ | 2,39 |
| OTU86 | Unclassified *Alphaproteobacteria* | 0,58 | $3,93 \times 10^{-9}$ | 2,08 |
| OTU58 | Unclassified *Rhodobacteraceae* | 0,56 | $1,69 \times 10^{-8}$ | 1,79 |
| OTU135 | Unclassified *Deltaproteobacteria* | 0,57 | $1,12 \times 10^{-8}$ | 1,42 |
| OTU96 | *Halioglobus* | 0,37 | $3,47 \times 10^{-4}$ | 1,30 |
| OTU136 | Uncultured *Saprospiraceae* | 0,59 | $1,93 \times 10^{-9}$ | 1,29 |
| OTU182 | *Microscilla* | 0,45 | $1,56 \times 10^{-5}$ | 1,26 |
| OTU172 | *Pseudohongiella* | 0,50 | $1,07 \times 10^{-6}$ | 1,11 |
| OTU202 | Unclassified *Rickettsiales* | 0,40 | $1,07 \times 10^{-4}$ | 1,06 |
| OTU143 | *Nisaea* | 0,55 | $4,50 \times 10^{-8}$ | 0,88 |
| OTU228 | Uncultured *Legionellaceae* | 0,43 | $3,46 \times 10^{-5}$ | 0,88 |
| OTU168 | Uncultured *Saprospiraceae* | 0,51 | $5,05 \times 10^{-7}$ | 0,84 |
| OTU241 | *Taeseokella* | 0,50 | $1,08 \times 10^{-6}$ | 0,62 |
| OTU288 | Unclassified *Babeliales* | 0,39 | $1,53 \times 10^{-4}$ | 0,62 |
| OTU296 | *Halioglobus* | 0,51 | $4,92 \times 10^{-7}$ | 0,36 |
| OTU306 | *Thalassobaculum* | 0,49 | $1,38 \times 10^{-6}$ | 0,31 |
| OTU445 | Uncultured *Legionellaceae* | 0,42 | $5,44 \times 10^{-5}$ | 0,31 |
| OTU375 | *Sneathiella* | 0,36 | $5,76 \times 10^{-4}$ | 0,27 |
| OTU371 | Unclassified *Rubinisphaeraceae* | 0,47 | $3,65 \times 10^{-6}$ | 0,27 |
| OTU75 | *Erythrobacter* | 0,37 | $3,89 \times 10^{-4}$ | 0,26 |
| OTU583 | Unclassified *Gammaproteobacteria* | 0,41 | $8,01 \times 10^{-5}$ | 0,22 |
| OTU293 | *Candidatus Peregrinibacteria* | 0,37 | $4,06 \times 10^{-4}$ | 0,22 |
| OTU305 | Unclassified *Gammaproteobacteria* | 0,39 | $1,80 \times 10^{-4}$ | 0,18 |
| OTU323 | *Thalassobaculum* | 0,40 | $1,17 \times 10^{-4}$ | 0,13 |
| OTU469 | *Pelagicoccus* | 0,35 | $8,65 \times 10^{-4}$ | 0,13 |
| OTU847 | *Marinoscillum* | 0,35 | $8,91 \times 10^{-4}$ | 0,05 |
| OTU537 | *Balneola* | 0,49 | $1,38 \times 10^{-6}$ | 0,04 |

Supplementary Table 2 – Bray-Curtis dissimilarities between the microbiomes from the live and dry feed products and exchange waters. The Bray-Curtis dissimilarities have been split into the turnover component (i.e. differences in presence/absence of taxa) and the part that is related to abundance variation (i.e. differences in relative abundance of the present taxa).

| | **Dissimilarity within the batches of this group** | | |
|---|---|---|---|
| | **Bray-Curtis** | | |
| | **Total** | **Turnover** | **Abundance variation** |
| ***Artemia*** | 0.72 ± 0.17 | 0.0018 ± 0.0013 | 0.72 ± 0.17 |
| **Algae** | 0.78 ± 0.17 | 0.0030 ± 0.0034 | 0.77 ± 0.17 |
| **Water exchange** | 0.83 ± 0.14 | 0.0009 ± 0.0007 | 0.83 ± 0.14 |
| **Dry feed** | 0.75 ± 0.18 | 0.0047 ± 0.0041 | 0.74 ± 0.18 |

Supplementary Table 3 – Bacterial taxa that were detected as core members in the algal bioreactors. The range of relative and absolute abundances for each core OTU in the algal bioreactors is reported as a measure of the relative importance of this OTU. OTUs are sorted according to the maximum relative abundance.

| **OTU** | **Genus** | **Minimal relative abundance (%)** | **Maximal relative abundance (%)** | **Minimal absolute abundance (cells/mL)** | **Maximal absolute abundance (cells/mL)** |
|---|---|---|---|---|---|
| Otu00001 | *Phaeodactylibacter* | 0,00 | 59,28 | 0 | $5,26 \times 10^6$ |
| Otu00057 | *Marinobacter* | 0,00 | 39,68 | 0 | $1,03 \times 10^6$ |
| Otu00021 | *Winogradskyella* | 0,00 | 18,40 | 0 | $4,78 \times 10^5$ |
| Otu00002 | *Marivita* | 0,00 | 17,70 | 0 | $8,31 \times 10^5$ |
| Otu00016 | *Alteromonas* | 0,00 | 12,89 | 0 | $5,96 \times 10^5$ |
| Otu00023 | *Aestuariibacter* | 0,00 | 11,69 | 0 | $5,48 \times 10^5$ |
| Otu00061 | *Neptuniibacter* | 0,00 | 9,25 | 0 | $8,21 \times 10^5$ |
| Otu00149 | *Marinobacter* | 0,00 | 7,58 | 0 | $1,80 \times 10^5$ |
| Otu00045 | *Muricauda* | 0,00 | 7,52 | 0 | $3,53 \times 10^5$ |
| Otu00349 | *Roseicyclus* | 0,00 | 5,22 | 0 | $2,10 \times 10^5$ |
| Otu00043 | *Nautella* | 0,00 | 3,40 | 0 | $1,60 \times 10^5$ |
| Otu00014 | unclassified *Rhodobacteraceae* | 0,00 | 1,53 | 0 | $7,64 \times 10^4$ |
| Otu00077 | *Methylophaga* | 0,00 | 0,53 | 0 | $1,37 \times 10^4$ |
| Otu00259 | *Methylophaga* | 0,00 | 0,50 | 0 | $3,49 \times 10^4$ |
| Otu00075 | *Erythrobacter* | 0,04 | 0,49 | $1,22 \times 10^3$ | $3,50 \times 10^4$ |
| Otu01463 | *Pseudohongiella* | 0,00 | 0,13 | 0 | $3,87 \times 10^3$ |

Supplementary Table 4 – Bacterial taxa that were detected as core members in the *Artemia* storage tanks. The range of relative and absolute abundances for each core OTU in the *Artemia* storage tanks is reported as a measure of the relative importance of this OTU. OTUs are sorted according to the maximum relative abundance

| **OTU** | **Genus** | **Minimum relative abundance (%)** | **Maximum relative abundance (%)** | **Minimum absolute abundance (cells/mL)** | **Maximum absolute abundance (cells/mL)** |
|---|---|---|---|---|---|
| Otu00008 | unclassified *Leuconostocaceae* | 0,40 | 93,45 | $4{,}01 \times 10^3$ | $1{,}11 \times 10^7$ |
| Otu00018 | *Pseudomonas* | 0,00 | 44,36 | 0 | $1{,}60 \times 10^6$ |
| Otu00022 | *Exiguobacterium* | 0,00 | 25,23 | 0 | $2{,}17 \times 10^6$ |
| Otu00035 | *Pseudoalteromonas* | 0,09 | 24,97 | $1{,}58 \times 10^3$ | $1{,}80 \times 10^6$ |
| Otu00004 | *Donghicola* | 0,04 | 19,13 | $2{,}22 \times 10^3$ | $3{,}51 \times 10^5$ |
| Otu00028 | *Catenococcus* | 0,00 | 18,46 | 0 | $9{,}54 \times 10^5$ |
| Otu00053 | *Vibrio* | 0,00 | 14,32 | 0 | $7{,}40 \times 10^5$ |
| Otu00088 | *Pseudomonas* | 0,00 | 13,94 | 0 | $3{,}04 \times 10^5$ |
| Otu00094 | unclassified *Rhodobacteraceae* | 0,00 | 11,24 | 0 | $3{,}98 \times 10^5$ |
| Otu00049 | *Halomonas* | 0,13 | 9,64 | $2{,}23 \times 10^3$ | $8{,}91 \times 10^5$ |
| Otu00014 | unclassified *Rhodobacteraceae* | 0,04 | 6,54 | $4{,}44 \times 10^3$ | $2{,}37 \times 10^5$ |
| Otu00075 | *Erythrobacter* | 0,00 | 6,02 | 0 | $5{,}57 \times 10^5$ |
| Otu00032 | *Mesoflavibacter* | 0,00 | 5,69 | 0 | $2{,}20 \times 10^5$ |
| Otu00092 | JGI 0000069-P22 | 0,00 | 5,54 | 0 | $2{,}87 \times 10^5$ |
| Otu00091 | *Arcobacter* | 0,00 | 5,04 | 0 | $1{,}10 \times 10^5$ |
| Otu00044 | *Rhodovulum* | 0,00 | 4,79 | 0 | $1{,}73 \times 10^5$ |
| Otu00016 | *Alteromonas* | 0,00 | 4,60 | 0 | $9{,}82 \times 10^4$ |
| Otu00006 | unclassified *Rhodobacteraceae* | 0,22 | 4,52 | $4{,}01 \times 10^3$ | $1{,}65 \times 10^5$ |
| Otu00110 | *Vibrio* | 0,00 | 4,50 | 0 | $2{,}32 \times 10^5$ |
| Otu00118 | *Pseudomonas* | 0,04 | 3,51 | $7{,}42 \times 10^2$ | $2{,}53 \times 10^5$ |
| Otu00029 | *Vibrio* | 0,00 | 3,42 | 0 | $2{,}47 \times 10^5$ |
| Otu00138 | *Marinomonas* | 0,00 | 2,66 | 0 | $1{,}38 \times 10^5$ |
| Otu00108 | *Halomonas* | 0,00 | 2,61 | 0 | $1{,}86 \times 10^5$ |
| Otu00105 | *Hyphomonas* | 0,00 | 2,35 | 0 | $7{,}15 \times 10^4$ |
| Otu00043 | *Nautella* | 0,04 | 1,33 | $2{,}22 \times 10^3$ | $4{,}05 \times 10^4$ |
| Otu00225 | *Halomonas* | 0,00 | 0,78 | 0 | $7{,}26 \times 10^4$ |
| Otu00015 | unclassified *Rhodobacteraceae* | 0,00 | 0,56 | 0 | $3{,}48 \times 10^4$ |

Supplementary Table 5 – Bacterial taxa that were detected as core members in the exchange water. The range of relative and absolute abundances for each core OTU in the exchange water is reported as a measure of the relative importance of this OTU. OTUs are sorted according to the maximum relative abundance

| **OTU** | **Genus** | **Minimum relative abundance (%)** | **Maximum relative abundance (%)** | **Minimum absolute abundance (cells/mL)** | **Maximum absolute abundance (cells/mL)** |
|---|---|---|---|---|---|
| Otu00006 | unclassified *Rhodobacteraceae* | 0,01 | 0,21 | $3,65 \times 10^2$ | $8,90 \times 10^4$ |
| Otu00004 | *Donghicola* | 0,00 | 0,17 | $5,62 \times 10^1$ | $7,62 \times 10^4$ |
| Otu00061 | *Neptuniibacter* | 0,00 | 0,11 | $7,73 \times 10^1$ | $5,21 \times 10^4$ |
| Otu00015 | unclassified *Rhodobacteraceae* | 0,00 | 0,08 | 0 | $3,00 \times 10^4$ |
| Otu00077 | *Methylophaga* | 0,00 | 0,07 | 0 | $2,98 \times 10^4$ |
| Otu00002 | *Marivita* | 0,00 | 0,06 | $7,73 \times 10^1$ | $2,56 \times 10^4$ |
| Otu00058 | unclassified *Rhodobacteraceae* | 0,00 | 0,05 | $1,83 \times 10^2$ | $1,77 \times 10^4$ |
| Otu00290 | unclassified *Bacteria* | 0,00 | 0,05 | 0 | $7,66 \times 10^2$ |
| Otu00044 | *Rhodovulum* | 0,00 | 0,03 | $2,39 \times 10^2$ | $1,17 \times 10^4$ |
| Otu00014 | unclassified *Rhodobacteraceae* | 0,01 | 0,03 | $3,09 \times 10^2$ | $8,89 \times 10^3$ |
| Otu00105 | *Hyphomonas* | 0,00 | 0,03 | $5,99 \times 10^1$ | $1,60 \times 10^3$ |
| Otu00047 | *Gilvibacter* | 0,00 | 0,01 | 0 | $5,27 \times 10^3$ |
| Otu00093 | unclassified *Rhodobacteraceae* | 0,00 | 0,01 | 0 | $6,06 \times 10^2$ |
| Otu00261 | *Ponticaulis* | 0,00 | 0,01 | 0 | $4,63 \times 10^3$ |
| Otu00034 | *Tropicibacter* | 0,00 | 0,01 | 0 | $4,31 \times 10^3$ |
| Otu00086 | unclassified *Alphaproteobacteria* | 0,00 | 0,01 | 0 | $2,87 \times 10^3$ |

Supplementary Table 6 - Physicochemical parameters of the cultivation tanks. DO, pH and temperature were measured in one of the five replicate tanks, for ammonia and nitrite a pooled sample for all tanks was used.

| Day | $NH_3$+ $NH_4^+$ (ppm) | $NH_3$ (ppm) | $NH_4^+$ (ppm) | $NO_2^-$ (ppm) | pH (-) | DO (ppm) | T (°C) | Alkalinity (meq/L) |
|---|---|---|---|---|---|---|---|---|
| 2 | 0,08 | 0,01 | 0,07 | 0,11 | 8,17 | 5,89 | 32,19 | 102,00 |
| 4 | 0,64 | 0,05 | 0,59 | 0,13 | 8,05 | 6,00 | 32,13 | 119,00 |
| 7 | 2,43 | 0,19 | 2,24 | 0,22 | 8,06 | 5,82 | 32,03 | 136,00 |
| 9 | 3,43 | 0,20 | 3,23 | 0,35 | 7,94 | 5,39 | 31,34 | 136,00 |
| 11 | 3,41 | 0,16 | 3,25 | 0,42 | 7,86 | 5,28 | 31,26 | 136,00 |
| 13 | 5,02 | 0,22 | 4,80 | 0,34 | 7,82 | 5,71 | 31,18 | 136,00 |
| 15 | 5,85 | 0,25 | 5,60 | 0,51 | 7,84 | 5,72 | 30,27 | 136,00 |
| 17 | 5,88 | 0,23 | 5,65 | 0,70 | 7,80 | 5,26 | 30,34 | 153,00 |

#### Supplementary Figures

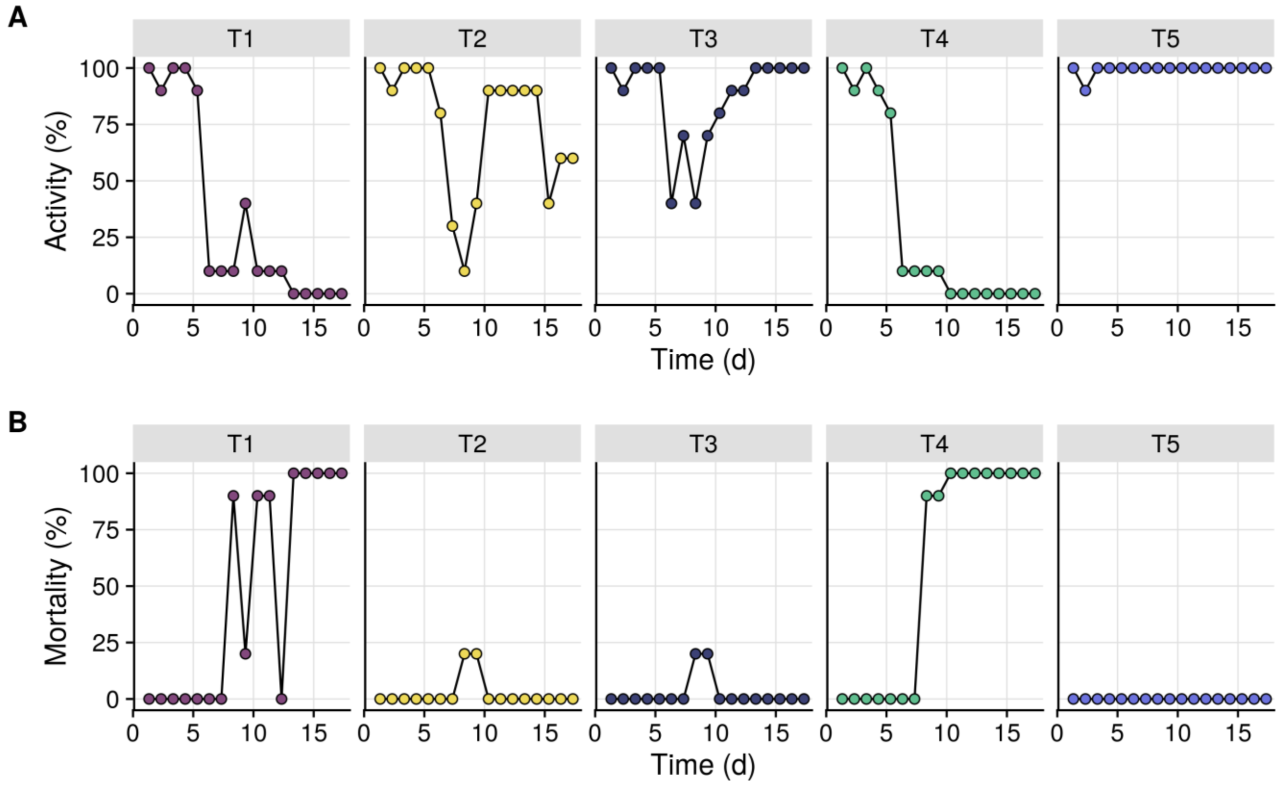


Supplementary Figure 1 – Daily estimations of larval activity (A) and cummulative mortality (B). These parameters are estimated based on visual inspection by the staff of the farm. The labels 'T1' to 'T5' refer to the five replicate tanks.

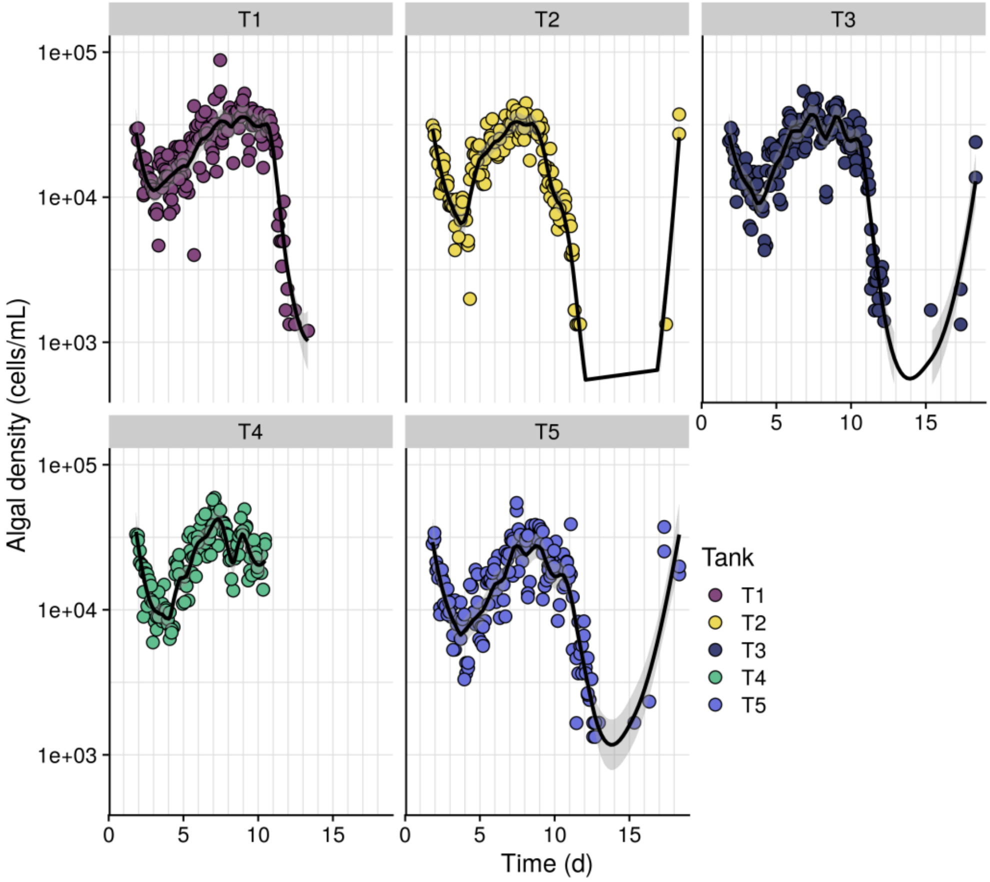


Supplementary Figure 2 – Algal abundances in the rearing water of the individual tanks over the cultivation period. Values below the detection limit (i.e. $10^3$ cells/mL) were removed from the dataset. The black lines indicate smoothing spline regressions and the shaded areas indicate corresponding 95% confidence intervals. The labels 'T1' to 'T5' refer to the five replicate tanks. It should be noted that at the end of the cultivation, algal densities were increased even though algae were not supplemented to the tank. The cause of this sudden increase is unknown.

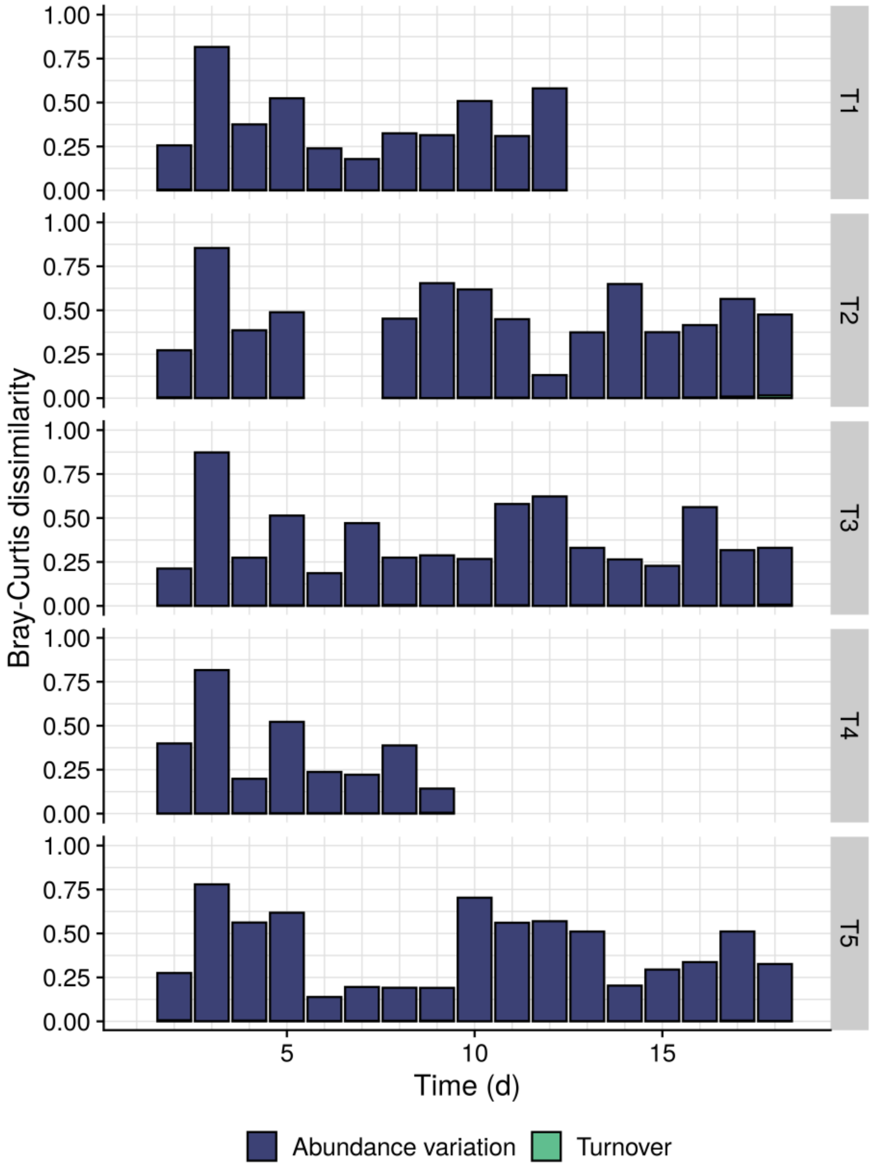


Supplementary Figure 3 – Bray-Curtis dissimilarities of the individual tanks over time, partitioned into the part that is caused by differences in absence/presence of taxa (i.e. turnover) and the part that is caused by differences in the relative abundances over the days (i.e. abundance variation). The value for a tank on day $i$ indicates the Bray-Curtis dissimilarity between the community in this tank on day $i$ as compared to the community in this tank on day $i$-1. Turnover was responsible for 0.71 % of the dissimilarity on average, and is therefore not visible on the graphs. For the two crashed tanks (i.e. T1 and T4), only the samples before the crash are included. For tank 2, there are two missing values because of a missing sample on day 6. The labels 'T1' to 'T5' refer to the five replicate tanks.

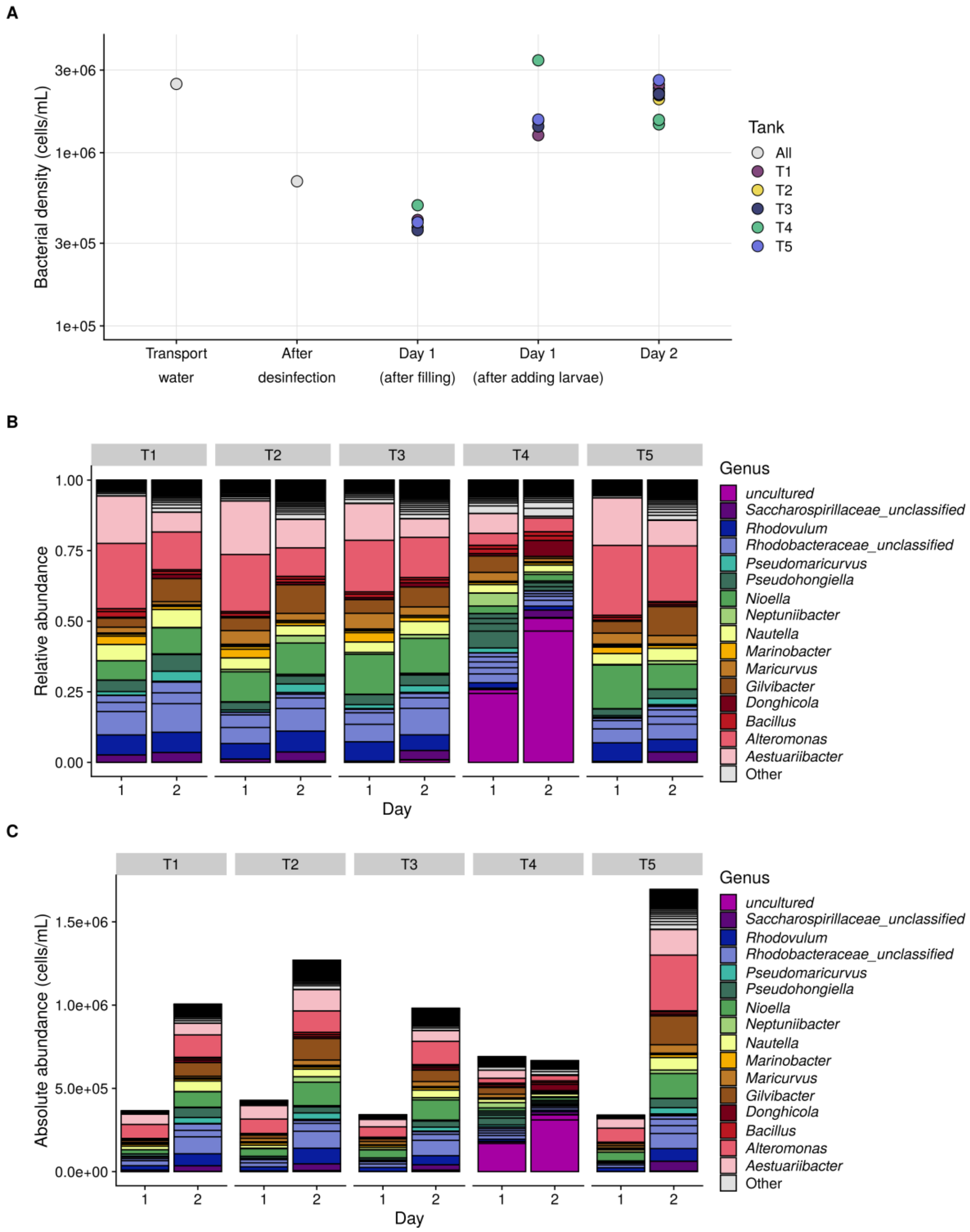


Supplementary Figure 4 – Bacterial abundance (A) and community composition (B, C) during the start-up and the first two days of the cultivation. The samples that are labelled “Day 1 (after filling)” are the rearing water microbiomes right after the tanks were filled with water, 8 hours before the addition of the nauplii. The transport water is the water in which the nauplii arrive at the farm, after which they are disinfected and distributed into the tanks (i.e. “Day 1 (after adding larvae)” samples). The sequencing of day 2 were taken the morning after (approx. 15 h later). The labels ‘T1’ to ‘T5’ refer to the five replicate tanks.

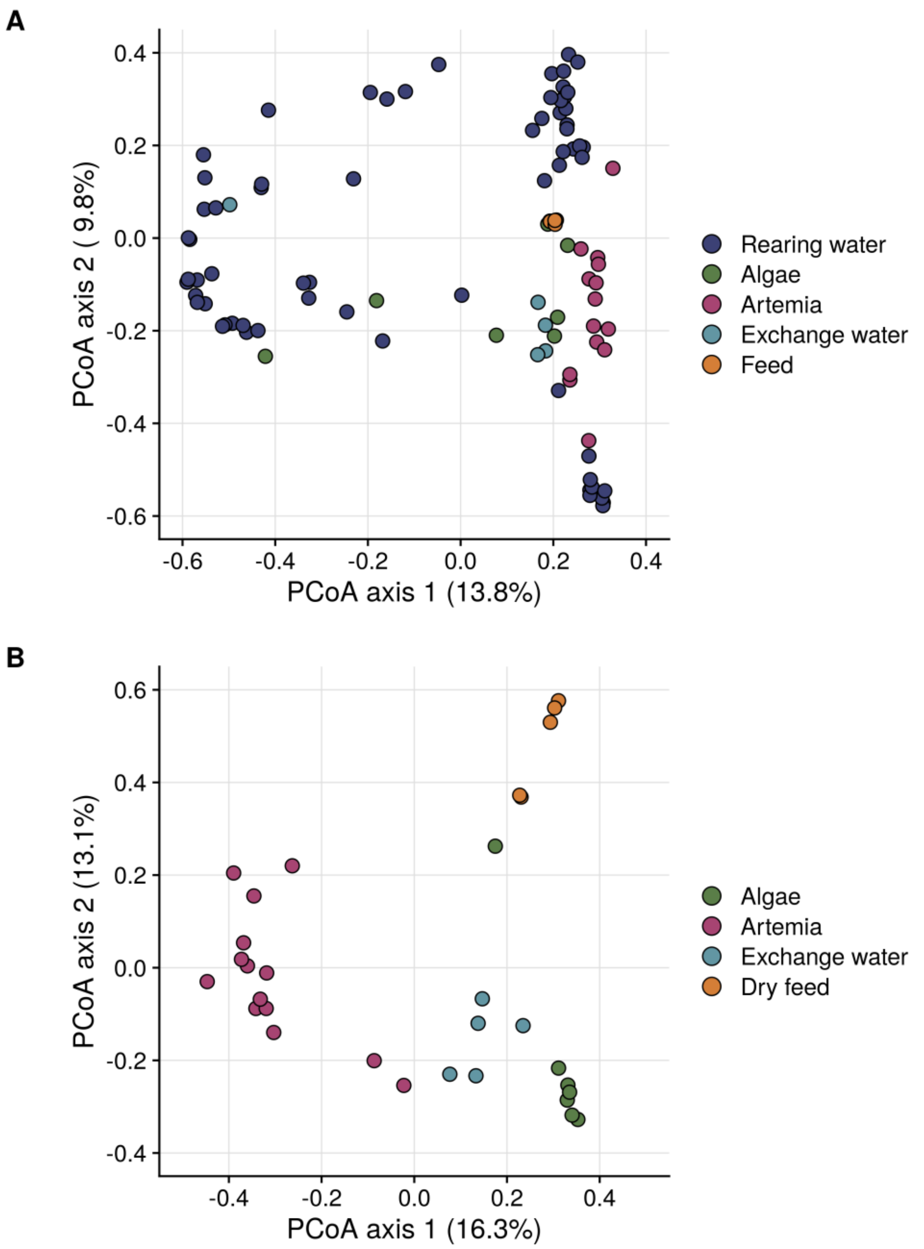


Supplementary Figure 5 - PCoA ordination of the Bray-Curtis dissimilarities of the rearing water and peripheral microbiomes (A) and peripheral microbiomes alone (B). Dots are coloured according to their origin. The microbiomes of the different types of peripheral microbiomes (i.e. algae, *Artemia*, dry feed and exchange water) were significantly different from one another. The variance explained by the source identity was 0.34 (p = 0.001, PERMANOVA). For the comparison between all peripheral microbiomes and the rearing water, no statistical test could be performed since the beta-dispersion of the different groups was significantly differing.

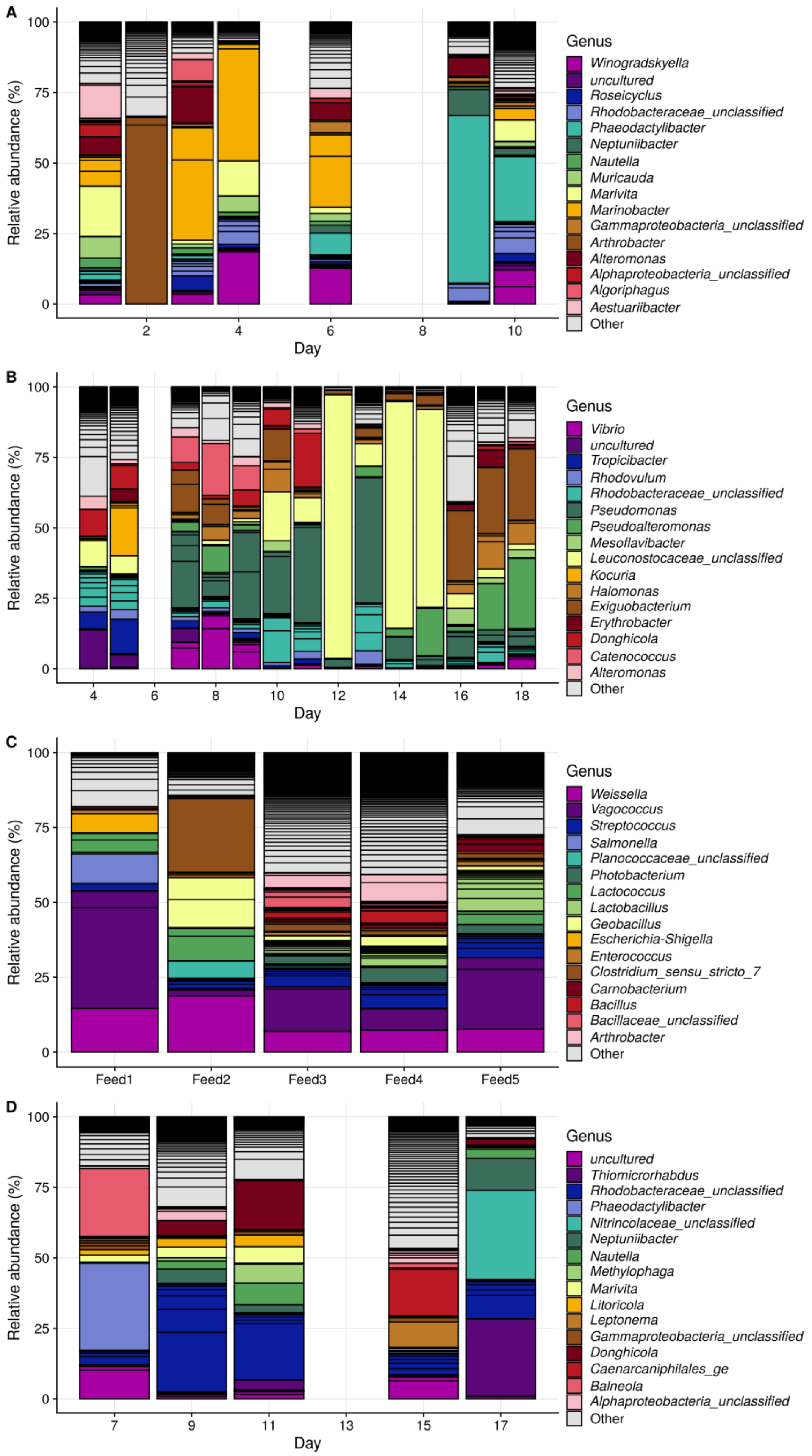


Supplementary Figure 6 – Community composition in the sources: algae (A), *Artemia* (B), dry feeds (C) and exchange waters (D). For each source, the OTUs belonging to the 16 genera with the highest overall abundance are coloured, all other genera are labelled as "Other".

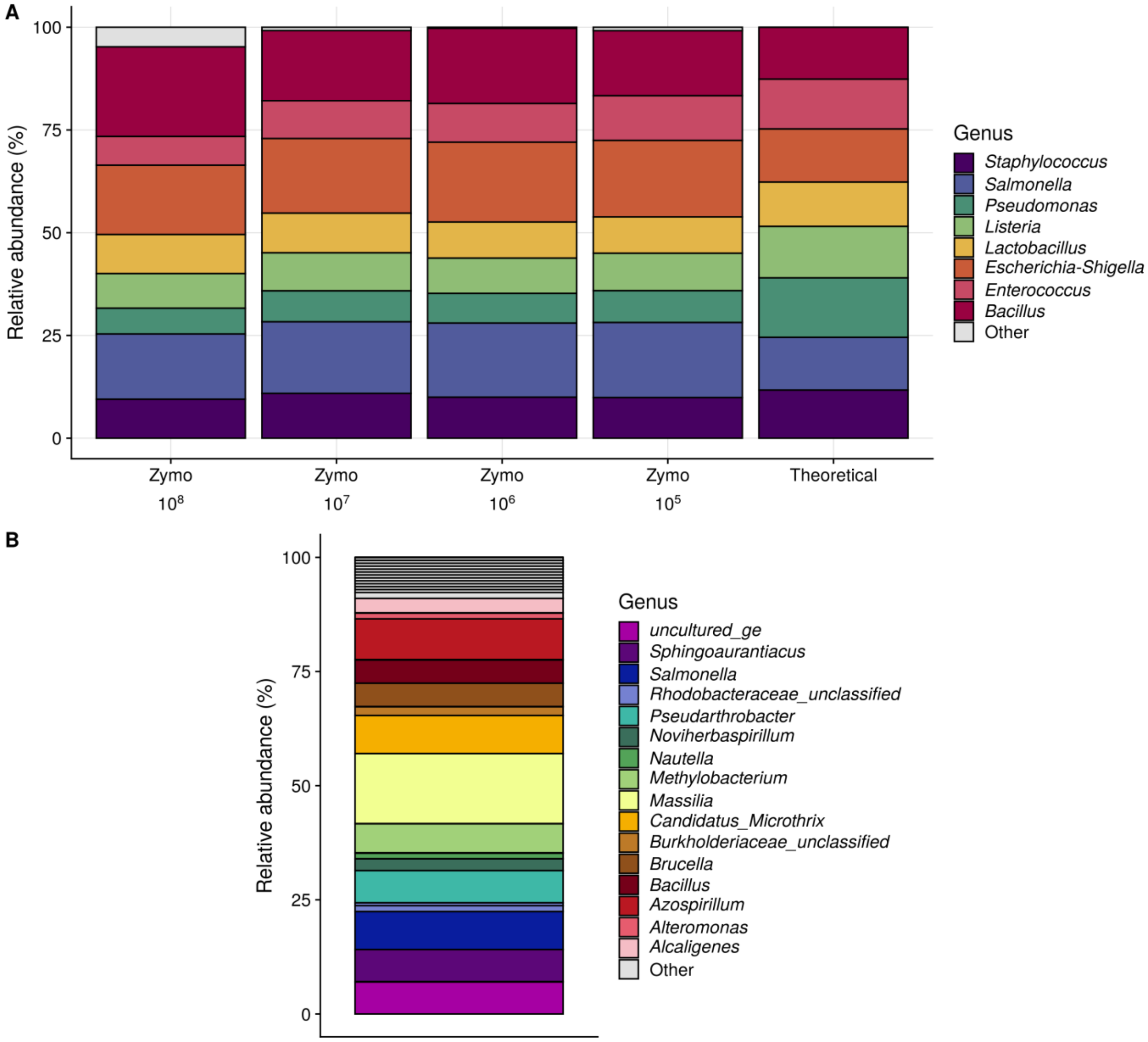


Supplementary Figure 7 – Quality controls for Illumina sequencing. (A) Community composition that was retrieved from the dilution series of the ZymoBIOMICS Microbial Community Standard (Zymo Research, USA). All contaminating OTUs are indicated as 'Other'. (B) Composition that was detected in an empty extraction with the ZymoBIOMICS DNA Microprep Kit (Zymo Research, USA). This sample was included in order to be able to control for potential laboratory or reagent contaminants. The OTUs belonging to the 16 genera with the highest overall abundance are coloured, all other genera are labelled as "Other".

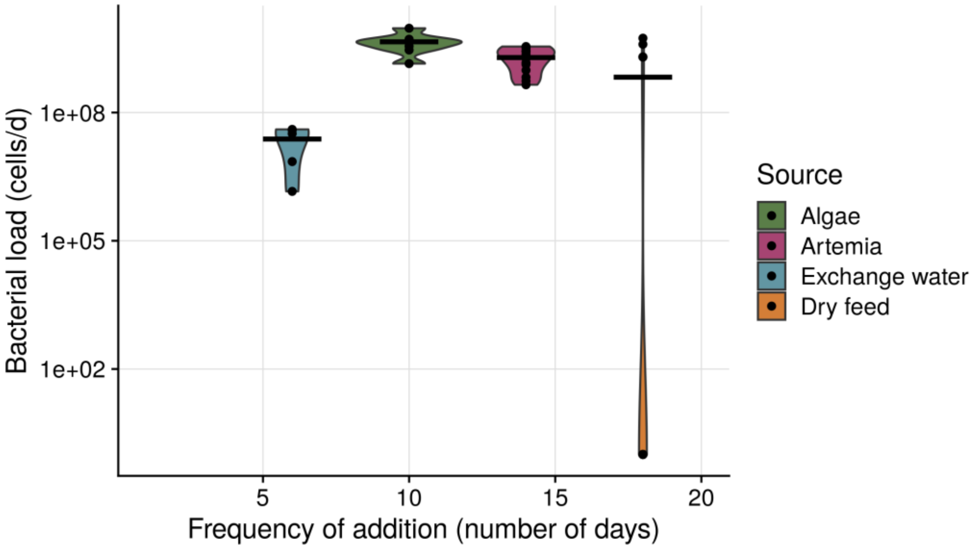


Supplementary Figure 8 – Frequency of addition and daily bacterial load towards the rearing water for each of the external sources. The horizontal lines indicate the average load over the entire cultivation, for each source. When taking into account the frequency of supplementation, dry feed caused a bacterial load of $7.47 \times 10^{8}$ cells towards the rearing water over the entire cultivation. For the algae this was $2.01 \times 10^{10}$ cells, for the *Artemia* $2.71 \times 10^{10}$ cells and for the exchange water $1.44 \times 10^{8}$ cells.

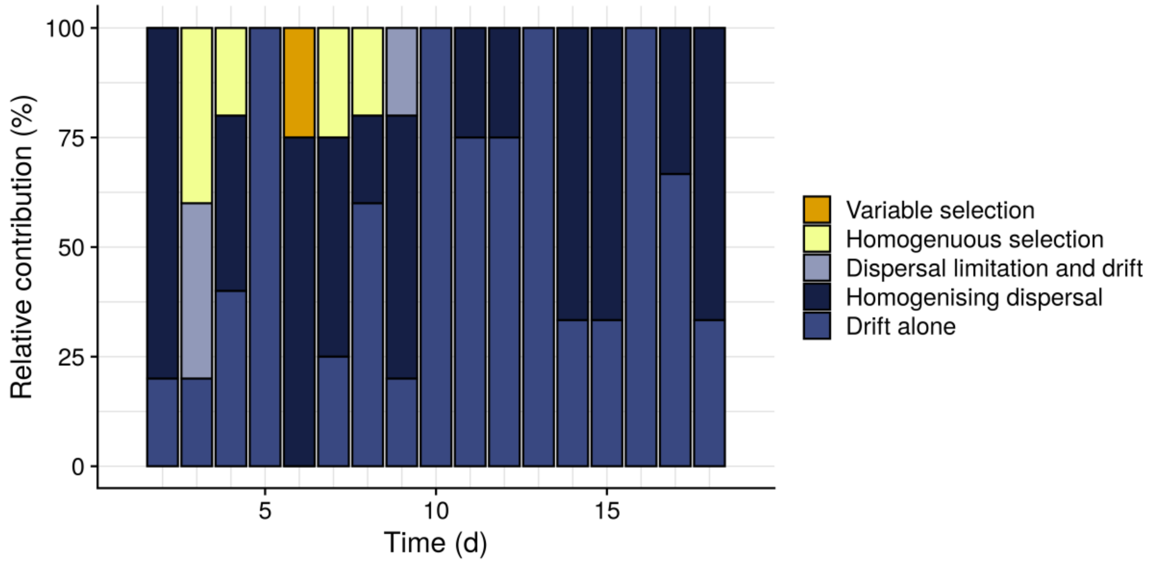


Supplementary Figure 9 – Distribution of the dominant community assembly processes in the rearing water over time, integrated over the replicate tanks. The bars on day $i$ indicate the relative contribution of the different processes for the assembly of the rearing water community of individual tanks from day $i$-1 to day $i$. The blue processes represent the stochastic assembly mechanisms, the yellow processes represent selective mechanisms. For the two crashed tanks (i.e. T1 and T4), only the samples before the crash are included.

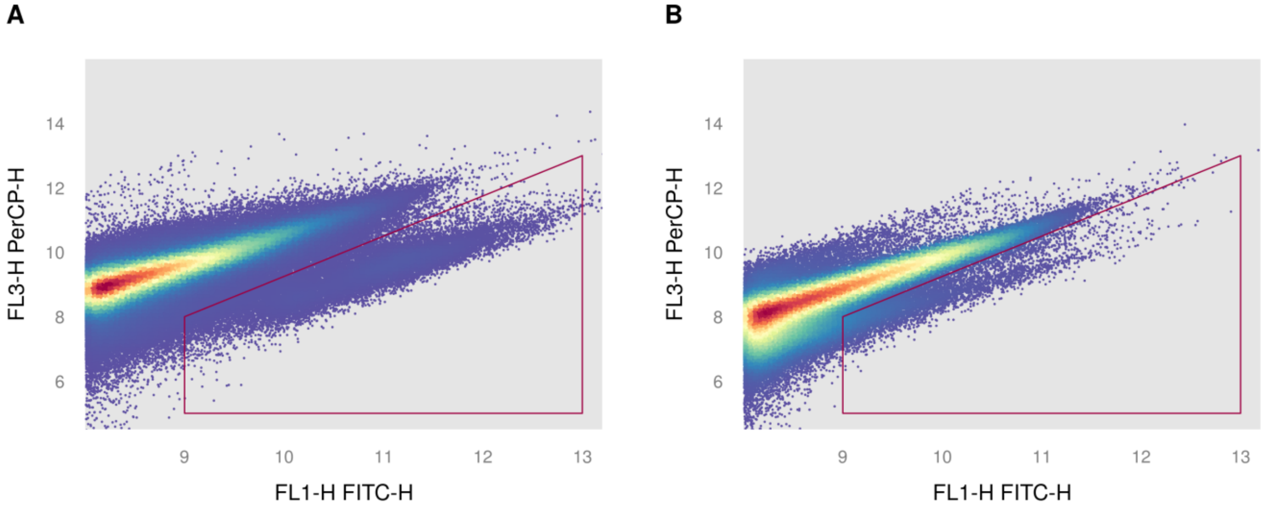


Supplementary Figure 10 – Illustration of the flow cytometry results for feed product 1 (A) and feed product 4 (B). The colour intensity is proportional to the log-scaled density of the events. For feed product 1 there is a clear separation between the cells and signals originating from the background. For feed product 4, events are observed in the region where cells are expected, but there is no clear separation between the cell population and the background.

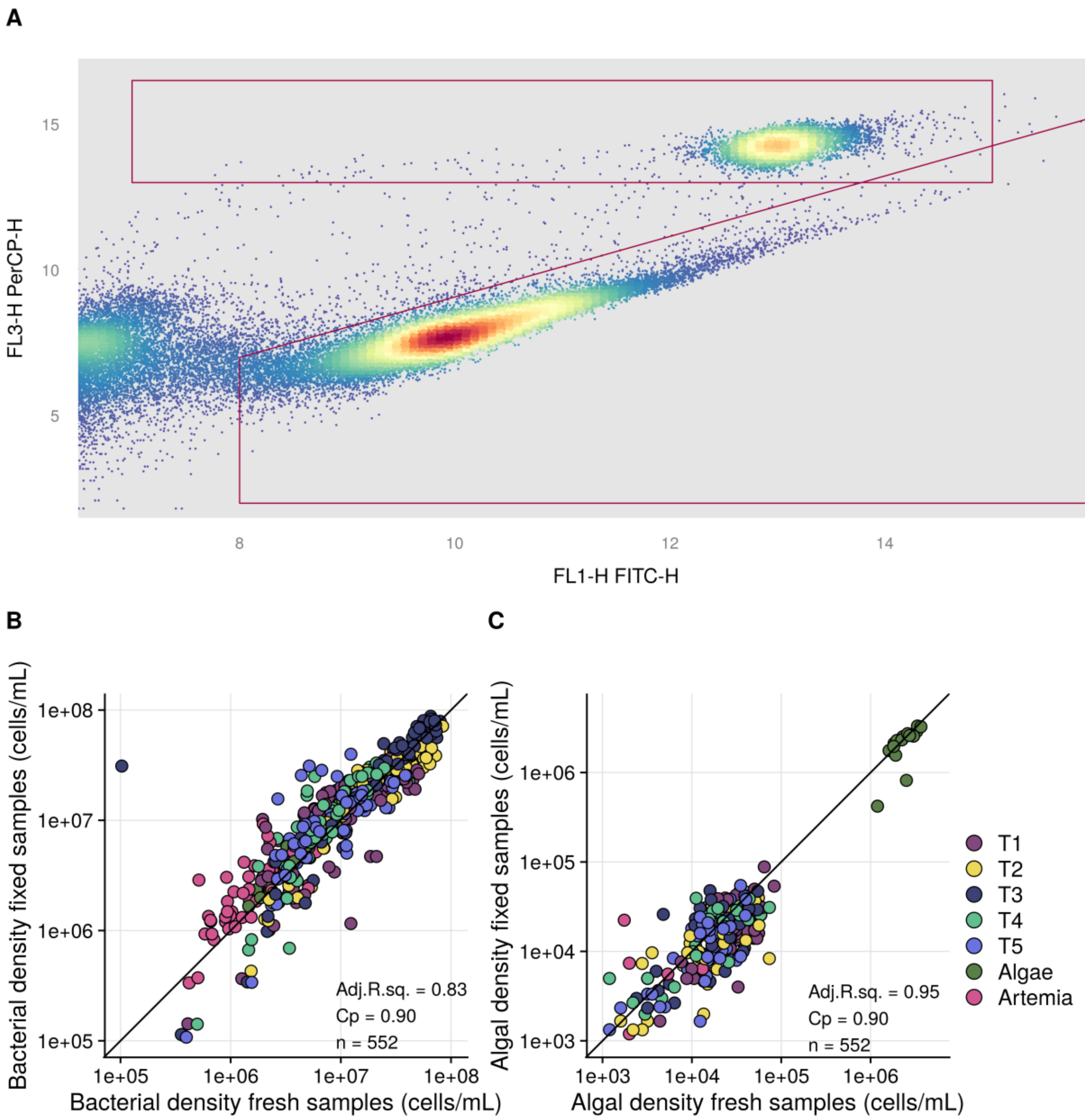


Supplementary Figure 11 – (A) Illustration of the gating strategy to separate bacterial and algal populations from the background signals. Gating was performed on the primary fluorescence channels. The colour intensity is proportional to the log-scaled density of the events. (B & C) Relationship between the bacterial (B) and algal (C) densities measured in the fresh samples on site, and the fixed samples off-site. The samples are coloured according to their origin (i.e. rearing water tanks or one of the live feed cultures). For both the bacterial and the algal densities, the fresh and fixed densities are highly correlated. The labels 'T1' to 'T5' refer to the five replicate tanks.
